# Computational Design of Periplasmic Binding Protein Biosensors Guided by Molecular Dynamics

**DOI:** 10.1101/2023.11.10.566541

**Authors:** Jack M. O’Shea, Annis Richardson, Peter Doerner, Christopher W. Wood

**Affiliations:** School of Biological Sciences, University of Edinburgh, Edinburgh, EH9 3FF, UK; School of Natural Sciences, Technical University of Munich, Center for Functional Protein Assemblies (CPA), Garching, 85748, Germany

## Abstract

Periplasmic binding proteins (PBPs) are bacterial proteins commonly used as scaffolds for substrate-detecting biosensors. In such biosensors, effector proteins, for example circularly permuted green fluorescent protein (cpGFP), are inserted into a PBP such that the effector protein’s output changes upon PBP-substate binding. The insertion site is often determined by comparison of PBP *apo*/*holo* crystal structures, but random insertion libraries have shown that this can miss the best sites. Here, we present a PBP biosensor design method based on residue contact analysis from molecular dynamics. This computational method identifies the best previously known insertion sites in the maltose binding PBP, and suggests further previously unknown sites. We experimentally characterise cpGFP insertions at these new sites, finding they too give functional biosensors. Our method is sufficiently flexible to both suggest insertion sites compatible with a variety of effector proteins and be applied to binding proteins beyond PBPs.

## Introduction

Protein-based biosensors are important tools for research in biochemistry and medicine. Being genetically encodable, they can monitor real-time *in vivo* conditions and can be optimised by directed evolution. The minimum requirement for a biosensor is that is possess an input-sensing “detector” domain and an output-generating “effector” domain. Modularity of detector and effector domains makes design of new sensors much simpler, which is why the periplasmic binding protein scheme of protein-based biosensors has been so successful.

Periplasmic binding proteins (PBPs) are a large family of bacterial proteins that scavenge and sense nutrients, and so are very well suited to detecting small molecules (Edwards, 2021; Nasu *et al*., 2021). They have evolved to bind a wide range of metabolically important ligands with high affinity and specificity, and new ligand specificities have been engineered by directed evolution (Guntas *et al*., 2005; Taylor *et al*., 2016) and computational methods (Tavares *et al*., 2019; Li *et al*., 2022). Upon substrate binding they undergo significant conformational change (figure 1A). For this reason their *apo* (without ligand) and *holo* (ligand-bound) states to be distinguished by receptors and thereby regulate many processes downstream in bacterial cells (Quiocho and Ledvina, 1996; Felder *et al*., 1999). Structurally, PBPs possess two distinct folded lobes that are spanned by flexible hinge loops, and upon substrate binding these lobes close around the substrate with a Venus flytrap- or hinge-like movement. In other words, when PBPs are *apo* (without ligand) they are most likely in an “open” conformation, and when they are *holo* (ligand-bound) they are in “closed” conformation (figure 1A).

**Figure 1.**
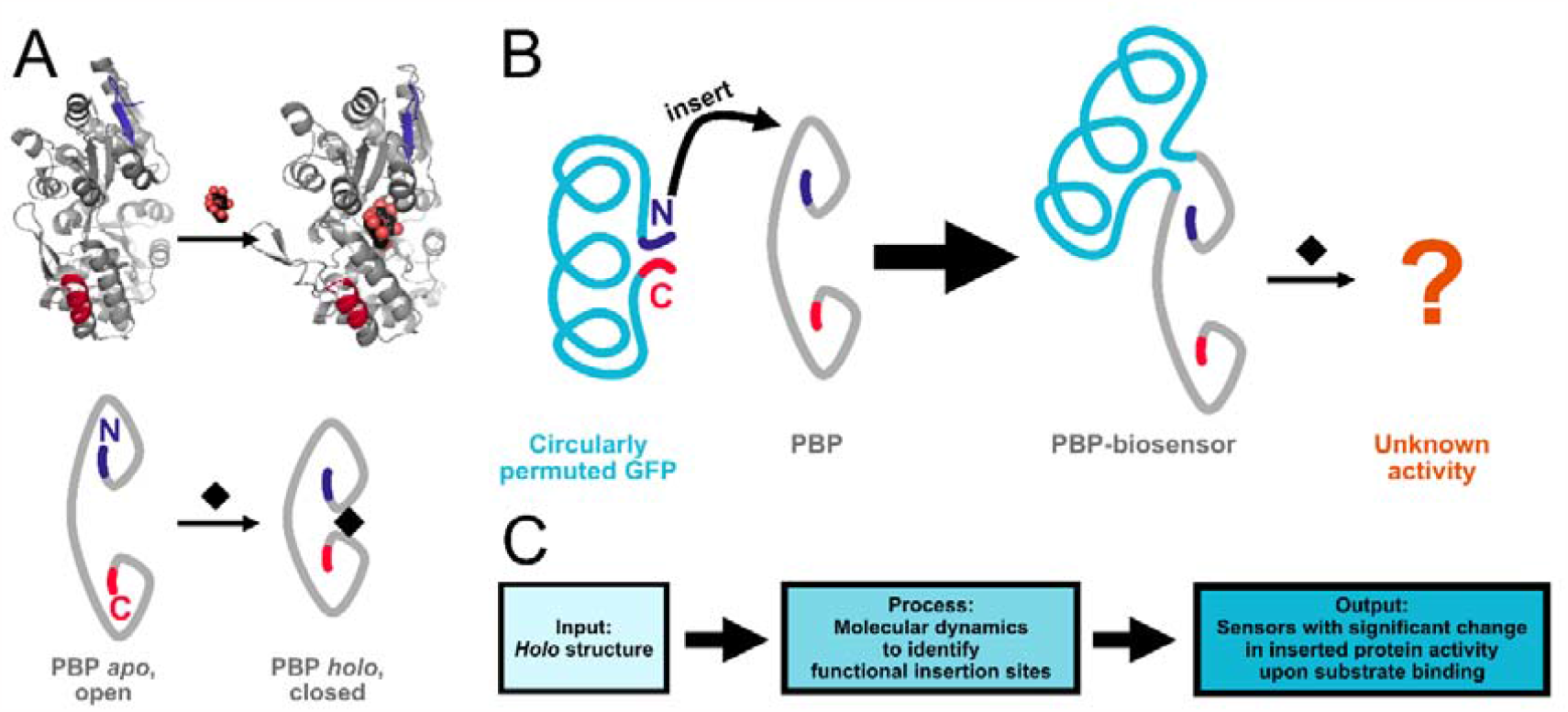
Schematics explaining design of PBP-biosensors. **A)** Crystal structures of *apo* (1ANF) and *holo* (1OMP) states of periplasmic binding proteins (PBPs), with schematic representations underneath. N and C termini are highlighted in blue and red respectively. **B)** Circularly permuted GFP has its N and C termini close together, so it can be inserted into PBPs to generate biosensors, but it is difficult to predict sensor activity from insertion site. **C)** Our method takes *holo* PBP structure as input, and using molecular dynamics simulations can provide insertion sites to generate functional biosensors.

In PBP biosensors, the PBP’s structural changes upon substrate binding are physically transmitted to a fused effector protein so as to measurably change the effector’s output. Sensors with outputs such as fluorescence (Alicea *et al*., 2011; Marvin *et al*., 2011, 2013; Nadler *et al*., 2016; Hu *et al*., 2018; Shivange *et al*., 2019; Borden *et al*., 2020; Jing *et al*., 2020), transcriptional repression (Younger *et al*., 2017, 2018), and enzymatic activity (Guntas and Ostermeier, 2004; Ribeiro *et al*., 2015, 2016; Choi, Xiong and Ostermeier, 2016) have been generated by inserting different effector proteins into PBP sequences, demonstrating the modularity of this approach. Typically, effectors are circularly permuted: their sequence is rearranged such that their N and C termini are very close to each other, ideal for insertion into the PBP sequence (Yu and Lutz, 2011; Ribeiro *et al*., 2019; Nasu *et al*., 2021). However, despite many examples of PBP biosensors, their design is impeded by our limited ability to identify sites in PBPs where effectors can be inserted to produce active sensors (figure 1B), which we term “the identification of functional insertion sites”.

Functional insertion sites were first rationally identified by the comparison of *apo* and *holo* crystal structures (Marvin *et al*., 2011). Marvin *et al*. showed that regions of greatest change in backbone dihedral torsion angles between the *apo*-open and *holo*-closed states of the maltose binding protein were associated with production of viable sensors upon insertion of circularly permuted GFP (cpGFP). This method has been successfully applied to other PBPs, either by referring to *apo*/*holo* crystal structures or by homology modelling of known successful insertion sites onto the new PBP (Alicea *et al*., 2011; Marvin *et al*., 2013, 2019). However, change in crystal structure backbone torsion angle can be an unreliable indicator of good insertion sites: sites of high change can be nonfunctional (Marvin *et al*., 2011) and sites of low change can be excellent (Nadler *et al*., 2016; Younger *et al*., 2018). Clearly crystal structure backbone torsion does not explain all factors necessary for functional insertion into PBPs. Alternatively, functional insertion sites have been identified by screens of random insertion libraries (Guntas and Ostermeier, 2004; Nadler *et al*., 2016; Ribeiro *et al*., 2016), but these methods are labour intensive, difficult to get full insertion coverage with, and do not explain why the identified insertion sites are functional.

In order to address these current shortcomings in PBP-based biosensor design, we reasoned that molecular dynamics (MD) could contain data that discern functional insertion sites better than crystal structure backbone torsion. We applied structural modelling, simulation, and analysis to the task (figure 1C) using the well-researched maltose binding protein (MBP) as a model subject. In simulations, requiring only the *holo* structure as a starting point, we captured MBP’s transition from closed to open state. We found that regions with substantial changes in residue contacts were correlated with the previously identified functional insertion sites, but the analysis also identified additional sites. From these additional sites, we generated new viable MBP sensors with a range of properties. We therefore propose that “change in residue contacts” is a more reliable indicator for functional insertion sites than “crystal structure backbone torsion”. This represents a significant advance in our understanding of the design parameters of PBP-based biosensors and demonstrates that computational analysis can greatly reduce barriers in the novel design of such sensors.

## Results and Discussion

Previous successful design of PBP biosensors has relied on inserting effector proteins into PBPs at regions of putative substrate-induced conformational change. We reasoned that functional insertion sites could be identified by using molecular dynamics simulations that capture such conformational change. As a test subject for this approach, we chose the maltose binding protein (MBP). Two previous studies provide screening data of large libraries of MBP with randomly inserted effector proteins. Nadler *et al*. randomly inserted a cpGPF, observing the greatest maltose-induced fluorescence changes from insertions at residues 169-171, followed by residues 355-348 (Nadler *et al*., 2016). Younger *et al*. inserted a zinc-finger transcription factor finding that insertion at position 335 produced the greatest maltose-induced change in transcription (Younger *et al*., 2018). Agreement in 335 as a good insertion site supports 355’s significance and shows that some insertion sites can be used with many different effectors. Therefore, the two regions of interest for our study are 169-171 and 335-348, and our aim was to find properties in molecular dynamics that distinguish these regions from all others.

We performed simulations of the substrate bound “closed” state using a crystal structure (PDB: 1ANF) as a starting point (Quiocho, Spurlino and Rodseth, 1997). 10 × 100 ns simulations were conducted (figure 2A, B). Open-state simulations were also performed from a starting point of 1ANF, except with maltose manually removed, and then simulations were run long enough for the protein to transition from the closed to the open state. This was a self-imposed restriction as we were aiming to produce a method that did not require crystal structures of both closed and open states (although both do exist for MBP). We ran 10 × 200 ns of these *apo* simulations, observing closed-to-open transition in 8/10 of them, visible as a distinct increase in RMSD (figure 2C, D). Representative endpoint poses of the *holo* and *apo* simulations clearly show the closed and open states respectively (figure 2E).

**Figure 2.**
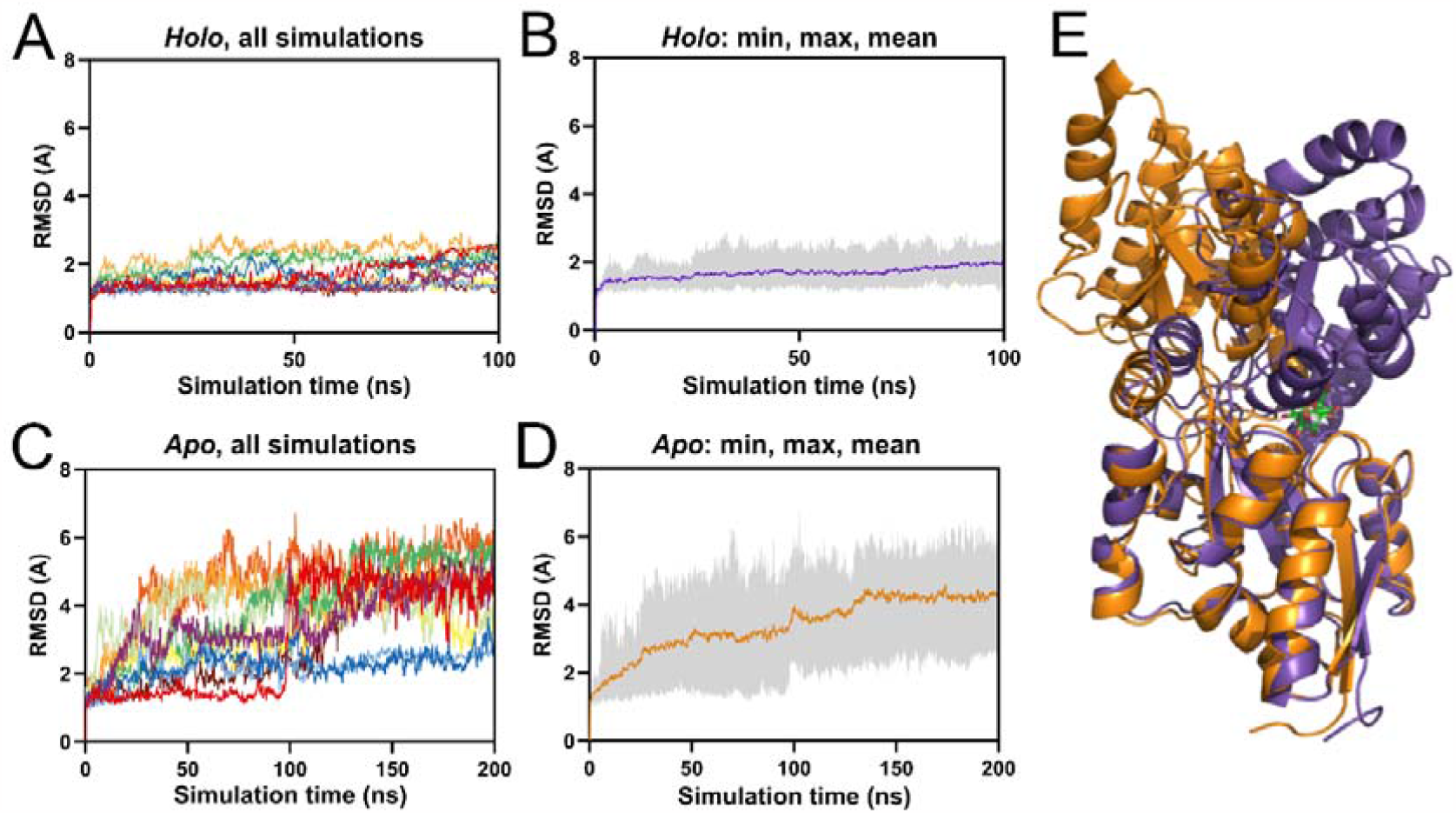
RMSD of molecular dynamics simulations of Maltose Binding Protein (MBP) in *apo* and *holo* forms. **A, B:** *Holo* simulations were generated from the starting point of the MBP/maltose complex crystal structure. **A)** Individual traces for 10x100ns MBP *holo* simulations. **B)** The mean, minimum, and maximum RMSD values for the MBP *holo* simulations. **C, D:** *A*po simulations were generated from the starting point of the MBP/maltose complex crystal structure with maltose removed. **C)** Individual traces for 10x200ns MBP apo simulations. **D)** The mean, minimum, and maximum RMSD values for the MBP apo simulations. **E)** Final poses of MBP in representative *holo*/closed (purple) and *apo*/open (orange) simulations, with maltose in green.

The first metrics we investigated were changes in backbone dihedral torsion (ΔDIHE-MD) and root mean squared fluctuation (ΔRMSF-MD) between closed and open simulations. For ΔDIHE at residue *i*, the dihedral angle between alpha-carbon atoms c-α_*i-1*_, c-α_*i*_, c-α_*i+1*_, and c-α_*i+2*_ is taken (Marvin *et al*., 2011). Between the MBP 1OMP (*holo*) and 1ANF (*apo*) crystal structures, residue 175 has the greatest ΔDIHE (Marvin *et al*., 2011), although Nadler and colleagues’ random insertion at residue 171 was found to produce a better sensor (Nadler *et al*., 2016). ΔRMSF was chosen to investigate regions of flexibility and conformational change within the protein. ΔDIHE and ΔRMSF were calculated for each residue for each of the final 50 ns of each closed-state simulation and for each of the simulations that reached the open-state, and the mean value in the open-state was subtracted from that of the closed. We compared these to ΔDIHE of the crystal structure (ΔDIHE-Crys). We hoped that at least one of ΔDIHE-MD and ΔRMSF-MD would highlight our regions of interest 169-171 and 335-348.

Unfortunately, ΔDIHE-MD and ΔRMSF were worse at highlighting our regions of interest than ΔDIHE-Crys (figure 3A, 3B, 3C). Residues in the region of interest (ROI) 169-171 have some of the largest ΔDIHE-Crys values, with 170 and 171 ranking 10^th^ and 11^th^. ΔDIHE-MD and ΔRMSF do not discern residues of this range at all. For the 335-348 range, residues did not stand apart for any measurements in MD or crystal structure. This shows that crystal structures can discern some good insertion sites (169-171) but not all (335-348). This initial analysis of the MD trajectory provided worse sites than using the static crystal structure alone, indicating that these metrics were not sensitive to the kinds of motions required to generate functional sensors.

**Figure 3:**
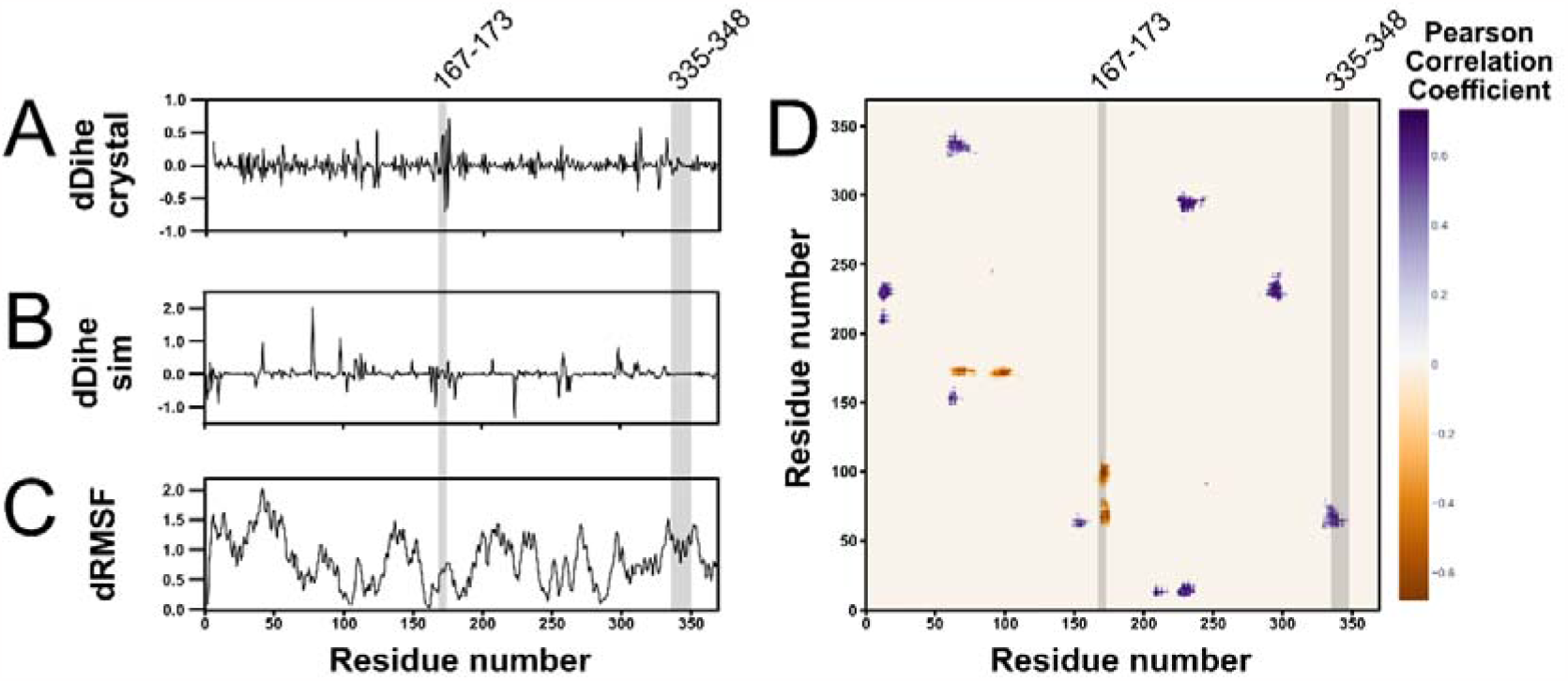
A comparison of metrics attempting to identify regions of MBP primary sequence known to be receptive to effector protein insertion with highlighted regions of interest. Across all figures, the residue ranges 167-173 and 335-348 are highlighted. 167-173 contains the region of interest 169-171, with flanking residues also highlighted for ease of visualisation. **A)** The change in backbone dihedral angle between *holo* and *apo* crystal structions (PDB 1ANF, 1OMP respectively). **B)** Change in backbone dihedral angle between *holo* and *apo* molecular dynamics simulations. Neither regions 167-173 nor 335-348 contain significant values for this metric. **C)** Difference in RMSF of each residue between *holo* and *apo* molecular dynamics simulations. **D)** Pearson correlation coefficient metric for all *apo* MBP simulations where the closed-to-open state transition was observed, filtered to only show peaks in the 99th percentile for mean, median, and maximum pixel value. Negative Pearson correlation coefficient indicates residues that become closer to each other throughout the closed-to-open transition, and positive values closer.

In order to identify more subtle structural dynamics, we applied the program CONAN (CONtact ANalysis) (Mercadante, Gräter and Daday, 2018) to our MD data. CONAN produces many metrics on the changes in interactions of a structure throughout a simulation. We submitted our *apo*-MBP simulations to CONAN, thereby providing it dynamic data on the close-to-open transition. Of interest to us was the Pearson Correlation Coefficient matrix output. This matrix would show us regions that experience changes in contacts between closed and open states, so may be involved in the changes in conformation that distinguish functional insertion sites.

We ran CONAN setting the contact cutoff as any residue atoms that came within 0.5 A and then remained within 0.8 A of each other. The resulting Pearson Correlation Coefficient matrix has positive values for residues that most interact early in the simulation (when MBP is closed) and negative values for residues that most interact late in the simulation (when MBP is open). To find the most significant hotspots of the Pearson Correlation Coefficient matrix, first an average-Pearson matrix was produced, with each pixel value being the mean of that pixel’s value across the Pearson matrices from each individual simulation (supplementary figure 1). Then pixels were clustered to define distinct peaks (positive values) and valleys (negative values). For each cluster, mean pixel value, median pixel value, and maximum absolute pixel value was calculated. Finally, peaks and valleys were filtered for those that were in the >99^th^ percentile of mean, median, and max absolute values simultaneously (peaks and valleys ranked separately).

In the resulting mean-filtered Pearson matrix (figure 3D), it is clear that our regions of interest 169-171 and 335-348 experience some of the most substantial changes in residue contacts during MBP opening in our MD. ROI 169-171 aligns with an area of negative Pearson space, indicating contacts that only occur in the open state. ROI 335-348 aligns with an area of positive Pearson space, indicating contacts that only occur in the closed state. The Pearson matrix calculated from the comparison of the MBP closed and open crystal structures (supplementary figure 2) has too much noise to discern any ROIs, demonstrating the necessity of MD-generated data. These data provide the first unifying known feature of ROIs 169-171 and 335-348: changes in contacting residues between closed and open state.

It is particularly of note that this metric captures ROI 335-348, which is not identified by comparison of crystal structures (figure 3A, B, C) but is identified by all random insertion studies to date (Nadler *et al*., 2016; Younger *et al*., 2018). These studies randomly inserted cpGFP and a zinc-finger transcription factor respectively and both identified residue 335, showing that functional insertion sites can be agnostic to the effector placed there. Another rational insertion study inserted beta-lactamase at 164 to achieve maltose-dependent antibiotic resistance (Guntas and Ostermeier, 2004), which is at the edge of the 170-centred CONAN hotspot. Changes residue contacts seems therefore to be a consistently important factor for a variety of sensor outputs, namely fluorescence, transcription, and catalysis. From this perspective, we propose that Pearson hotspots therefore correspond to regions where an inserted effector will experience significant changes in residue environment upon substrate binding, and that these changes are likely to significantly modulate effector output.

To validate this rationale for identifying good MBP insertion sites, we used the Pearson data to guide the design of new maltose sensors. We decided to explore the areas of positive Pearson correlation highlighted in our mean-filtered matrix from residue 200 to 250 (figure 3D). This region is sparsely sampled by random insertions with the transposon-based method, and is not distinguished by crystal structure ΔDIHE, so could provide previously unknown sensors.

Sequences were generated for the insertion of cpGFP into MBP every three residues in clusters 205-217 and 224-233. Sensors will be denoted by “**MBP_i-cpGFP**”, where **i** is the residue index of the insertion. For linkers, we emulated the design of Nadler *et al*. (Nadler *et al*., 2016) (nucleotide and amino acid sequences in supplementary table 3). The resulting sequences were cloned into pET-28a backbone. For negative controls, cpGFP was inserted at regions of flat Pearson space: **MBP_131-cpGFP** and **MBP_195-cpGFP**. For positive controls, two very good sensors from Nadler *et al*. were reproduced: **MBP_170-cpGFP** and **MBP_348-cpGFP** (Nadler *et al*., 2016).

Plasmids for expression of these sensors were transformed into BL21 *E. coli* and induced with IPTG. Pellets were recovered and lysed. Lysates were mixed with a pseudocytosol to simulate eukaryotic cellular conditions. The pseudocytosol was supplemented with either 0 mM, 1 mM, or 100 mM maltose, and the samples were tested for cpGFP fluorescence on a plate reader. Sensors were assayed by two metrics: raw fluorescence (arbitrary units) and ΔF/F_0_ (where ΔF is the difference in fluorescence between 0 mM maltose and the given concentration, and F_0_ is the fluorescence at 0 mM). ΔF/F_0_ is a key metric in fluorescent sensor design, with an absolute value of > 0.25 being considered a successful design (Nasu *et al*., 2021). The absolute ΔF/F_0_ values for our negative controls **MBP_131-cpGFP** and **MBP_195-cpGFP** did not exceed 0.25.

By this standard, and when compared to previously discovered sensors, our method has generated several new viable maltose sensors (figure 4, supplementary table 1, 2). For example, **MBP_233-cpGFP** had lower raw fluorescence than those of Nadler’s **MBP_170-cpGFP** and **MBP_348-cpGFP**, but in terms of ΔF/F0 **MBP_233-cpGFP** exhibited higher values under our experimental conditions. At concentrations of 1 and 100 mM, the ΔF/F0 values of **MBP_233-cpGFP** were 0.729 and 0.741, respectively, while for **MBP_170-cpGFP** they were 0.495 and 0.523, and for **MBP_348-cpGFP** they were 0.445 and 0.650. By the “>0.25” standard, **MBP_233-cpGFP** is certainly a viable sensor.

**Figure 4.**
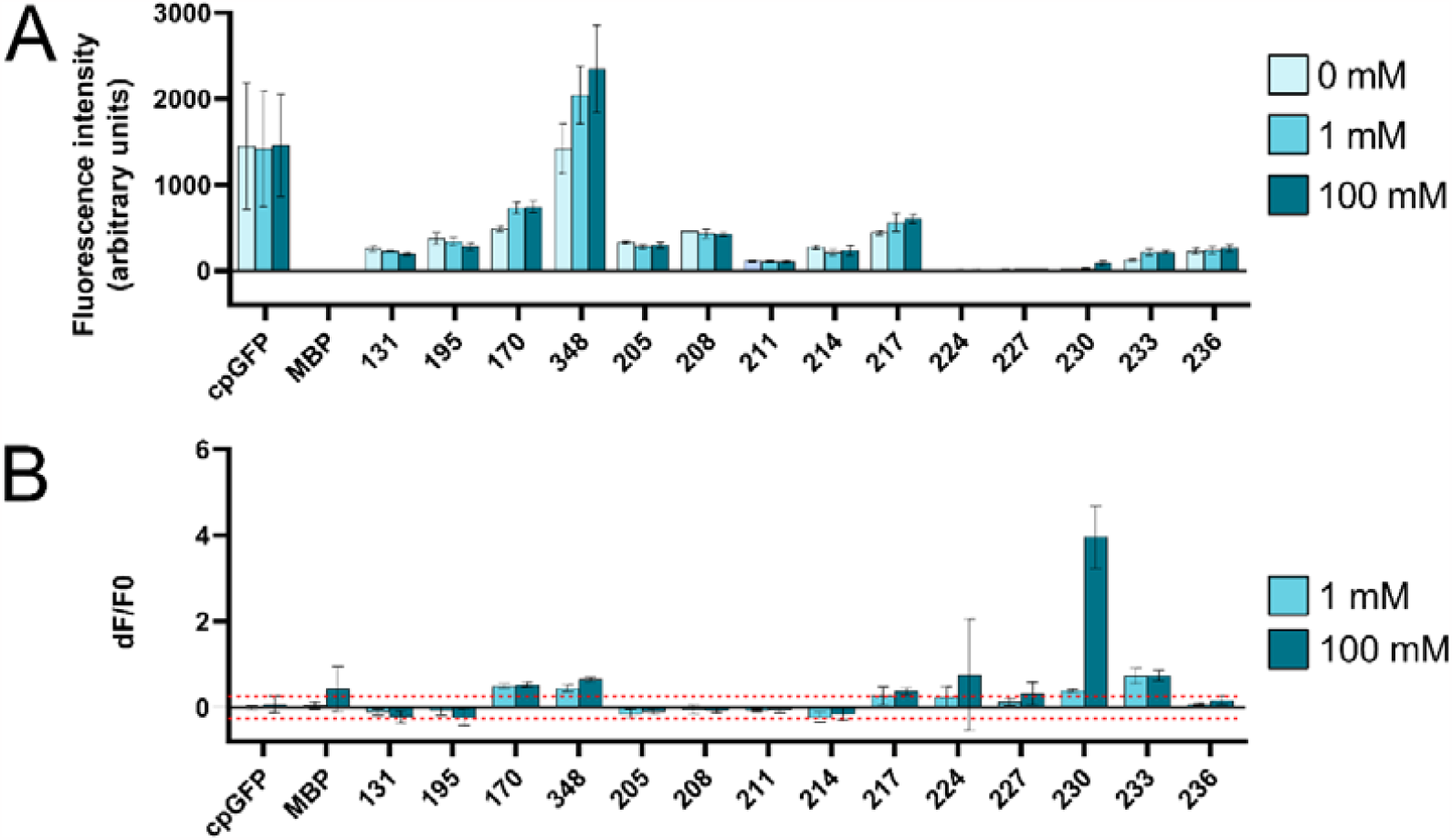
comparison of fluorescence of designed sensors at varied maltose concentrations. On the left of the axes are the control conditions. cpGFP: the fluorescent protein alone. MBP: the Maltose Binding Protein alone. 131 and 195: cpGFP inserted at MBP’s positions 131 and 195, which are both in areas of flat Pearson space so are predicted to generate poor quality sensors. 170 and 348: cpGFP inserted at MBP’s positions 170 and 348, both very strong sensors previously identified by Nadler et al. (Nadler *et al*., 2016). Presented are the means and standard deviations of 3 biological repeats per condition. Tables for all values are in supplement. **A**) Raw fluorescence of sensors at 0, 1, and 100 mM maltose. **B)** Relative change in fluorescence from 0 mM maltose to 1 or 100 mM, calculated by (F_0_ - F_X_) / F_0_. 0.25 and -20.5 are indicated with dashed lines.

**MBP_230-cpGFP** is another sensor of note, with low raw fluorescence but by far the greatest ΔF/F_0_. It exhibited very little fluorescence at 0 and 1 mM and was only slightly brighter at 100 mM, with raw measured values of 22, 31, and 72 respectively. However, this small change resulted in a 100 mM ΔF/F_0_ of 3.954, by far the largest of those we measured. **MBP_230-cpGFP’**s fluorescence uptick at 100 mM was extremely consistent between experimental repeats, not a result of small fluctuations around a small value, making it another viable sensor, particularly for use in conditions of low background fluorescence. **MBP_227-cpGFP** also achieved ΔF/F_0_ > 0.25 for 100 mM, and **MBP_217-cpGFP** did so at 1 and 100 mM, totalling 4 identified viable sensors.

Not all designs were successful. **MBP_224-cpGFP** and **MBP_227-cpGFP** had no fluorescence over all conditions (the apparent large ΔF/F0 for **MBP_224-cpGFP** at 100 mM is an artefact of small fluctuations). Insertions from 205-214 did not provide viable sensors, with small if any ΔF/F_0_ values. Clearly yet further factors than Pearson hotspots determine whether an insertion is functional or not. That said, the identification of 4 new viable sensors with a variety of profiles generated by our method is very encouraging, especially since we purposefully restricted our search to regions that were not well sampled by previous studies. Although our designs are less bright than those of previous studies, brightness is one feature that can be relatively easily improved by directed evolution.

Modulation of effector domain output upon substrate binding is central to biosensor design. Early PBP-biosensor studies inserted effectors at regions of high backbone dihedral angle change (high ΔDIHE), and attributed output modulation to direct transmission of backbone conformational changes from detector to effector. However, random insertion studies have shown that some insertions at sites of low ΔDIHE give substantial substrate-induced output changes, showing that ΔDIHE the insertion sites that best modulate effector output are sometimes overlooked by this metric. Here we use Pearson Correlation Coefficient of residue contacts in MD simulations to show that regions of significant contact change during PBP opening correlate well with the known best insertion sites. We then designed new sensors from insertions in regions of high Pearson Correlation Coefficient and low ΔDIHE, validating this as a new method to generate PBP-biosensors.

In this work we have demonstrated that Pearson hotspots correlate with known good insertion sites from more regions of primary sequence better than changes in dihedral angles of crystal structures or MD-derived poses. The Pearson Correlation Coefficient matrix can be accessed from only the *holo* crystal structure, and could even be accessed for PBPs with no known crystal structure thanks to structural prediction algorithms such as AlphaFold2, which are known to mostly predict *holo* states of binding proteins (Saldaño *et al*., 2022). A wider variety of candidate insertion sites will help with the successful insertions of a wider variety of effectors such as transcription factors and enzymes (Guntas and Ostermeier, 2004; Younger *et al*., 2018). It would also be interesting to see this method applied to other sensor proteins with similar hinge-like binding dynamics, such as the *de novo* hinge proteins recently developed in the Baker lab (Praetorius *et al*., 2023). Finally, the most immediate impact of this work is the forging of a new, computational avenue for PBP-biosensor discovery that can be performed at a much greater scale than current wet-lab methods for minimal labour and cost.

## Methods

### MD simulations

Simulations were set up using AmberTools and performed using the Python package OpenMM (Eastman *et al*., 2013). Nonbonding interactions were modelled by PME, with cut-off at 1 nm. Simulations were run at 1 bar with a Monte Carlo barostat and 300 K temperature was maintained by the Langevin integrator with frictional constant 1 ps^-1^. Hydrogen bond length constrains were applied. A timestep of 2 fs was used. Code for running these simulations can be found at https://github.com/wells-wood-research/oshea-j-wood-c-pbp-design-2023.

### RMSD, RMSF, dihedral torsion

RMSD and RMSF calculations were performed using the Python package MDAnalysis (Gowers *et al*., 2016). Dihedral torsion angles were calculated using the Python package MDTraj (McGibbon *et al*., 2015). Code for calculating the change in RMSF and dihedral torsion angles can be found at https://github.com/wells-wood-research/oshea-j-wood-c-pbp-design-2023.

### CONtact ANalysis (CONAN)

CONAN was applied to the 8/10 *apo* simulations where MBP opening occurred. We defined a “contact” as any residue atoms that came within 0.5 A and then remained within 0.8 A of each other, as this is well within the limit to exclude water (which has a radius of 1.9 A). The input file and outputs relevant to this work can be found at https://github.com/wells-wood-research/oshea-j-wood-c-pbp-design-2023. For each contact in the Pearson Correlation Coefficient matrix, the pixel’s mean value across the 8 simulations analysed was taken. These means formed the matrix that was subjected to pixel clustering, the code for clustering was adapted from that of Galloway *et al*. (2021) to differentiate between peaks and valleys. For each cluster, mean pixel value, median pixel value, and maximum absolute pixel value was calculated. Peaks and valleys were then filtered for those that were in the >99^th^ percentile of all of these values (peaks and valleys ranked separately).

### Design of constructs for expression of sensors

For a given insertion point, linker design followed that of Nadler *et al*. (2016). N-terminal linkers were *res*^*i+1*^-Ala-Ser, where *res*^*i+1*^ is the residue after the inserted residue. C-terminal linkers were Ala-Ser-*res*^*i*^, where *res*^*i*^ is the residue that was inserted at. The amino acid sequence of cpGFP flanked by the linkers was inserted into the amino acid sequence for MBP. The amino acid sequence was codon optimised for expression in *Escherichia coli* using the Twist Bioscience sequence input interface. Optimised sequences were cloned into the pET28a backbone at the SacI_HindIII restriction sites.

### Cloning of constructs

Primers for cloning and sequencing were ordered from IDT. Fragments were amplified using Phusion polymerase (NEB), run on 1% agarose gels and recovered using GeneJET Plasmid miniprep kit (ThermoFisher). Recovered fragments were assembled using NEBuilder kit (NEB). Assembled plasmids were sequenced by the Medical Research Council Protein Phosphorylation and Ubiquitylation Unit’s DNA sequencing service at the University of Dundee.

### Sensor expression

Plasmids were transformed into BL21-alpha cells (NEB) and plated overnight at 37 □C on kanamycin LB plates (50 μg/mL). The next day, colonies were picked into 2 mL of TB supplemented with kanamycin (50 μg/mL), and incubated at 37 □C, 190 rpm for 20 hours. 500 uL of these cultures were then used to inoculate 25 mL of LB supplemented with kanamycin (50 μg/mL), and incubated at 37 □C, 240 rpm for 1.5 hours. Cultures were then supplemented with 1 mM IPTG, and further incubated at 25 □C, 240 rpm for 20 hours.

### Fluorescent assay

After expression, cultures were centrifuged at 4000 G, 4 □C for 15 minutes. The pellets were recovered and weighed, then resuspended in 1 uL/mg of lysis buffer (1x BugBuster (Novagen), 20 mM Tris-HCl, 150 mM K-gluconate, pH 7.5 (Mukherjee *et al*., 2015)). Lysis was performed with shaking at room temperature, 1600 rpm, 30 minutes. Lysates were spun down at 16000 G for 15 minutes and the supernatant was recovered. 20 uL of supernatant was transferred to wells in the 96 well black/clear bottom plate (ThermoFisher). 200 uL of pseudocytosol (100 mM K-gluconate, 30 mM NaCl, 25 mM MES, 25 mM HEPES, 40% sorbitol, 1mg/mL bovine serum albumin, pH 7.5 (Messerli and Robinson, 1998; Feijó *et al*., 1999; Mukherjee *et al*., 2015)) supplemented with 0, 1, or 100 mM maltose. Plates were immediately imaged on an Infinite M200PRO (Tecan) plate reader, with excitation/emission of 485/515nm, with gain set to 40. In each of the three biological repeats, three technical repeats were performed for each condition and from these the mean was calculated. The mean value of the background negative control pET28a was subtracted from the positive controls and experimental conditions. From this background-subtracted mean value, ΔF/F_0_ values were calculated, where ΔF = F_X_ – F_0_, where X = 1 or 100. For the MBP negative control, 5 μg of MBP in 5 μL (RayBiotech) was mixed with 15 μL of lysis buffer and 200 μL of pseudocytosol with 0, 1, or 100 mM maltose.

## Acknowledgements

This work was supported by a BBSRC Engineering Biology Breakthrough Award (BB/W013320/1). Thank you to Dr. Frank Machin of the Doerner lab for direction to literature and Jonathan Lecoy of the Richardson lab for help using their lab space. Thank you to the teams at the Edinburgh Genome Foundry and the Edinburgh Protein Production Facility.

## References

Alicea, I. et al. (2011) ‘Structure of the Escherichia coli Phosphonate Binding Protein PhnD and Rationally Optimized Phosphonate Biosensors’, Journal of Molecular Biology, 414(3), pp. 356–369. Available at: 10.1016/j.jmb.2011.09.047.

Borden, P.M. et al. (2020) ‘A Fast Genetically Encoded Fluorescent Sensor for Faithful in vivo Acetylcholine Detection in Mice, Fish, Worms and Flies’. Rochester, NY. Available at: 10.2139/ssrn.3554080.

Choi, J.H., Xiong, T. and Ostermeier, M. (2016) ‘The interplay between effector binding and allostery in an engineered protein switch’, Protein Science, 25(9), pp. 1605–1616. Available at: 10.1002/pro.2962.

Eastman, P. et al. (2013) ‘OpenMM 4: A Reusable, Extensible, Hardware Independent Library for High Performance Molecular Simulation’, Journal of Chemical Theory and Computation, 9(1), pp. 461–469. Available at: 10.1021/ct300857j.

Edwards, K.A. (2021) ‘Periplasmic-binding protein-based biosensors and bioanalytical assay platforms: Advances, considerations, and strategies for optimal utility’, Talanta Open, 3, p. 100038. Available at: 10.1016/j.talo.2021.100038.

Feijó, J.A. et al. (1999) ‘Growing pollen tubes possess a constitutive alkaline band in the clear zone and a growth-dependent acidic tip’, The Journal of Cell Biology, 144(3), pp. 483–496. Available at: 10.1083/jcb.144.3.483.

Felder, C.B. et al. (1999) ‘The venus flytrap of periplasmic binding proteins: An ancient protein module present in multiple drug receptors’, AAPS PharmSci, 1(2), p. 2. Available at: 10.1208/ps010202.

Galloway, J.M. et al. (2021) ‘De Novo Designed Peptide and Protein Hairpins Self-Assemble into Sheets and Nanoparticles’, Small, 17(10), p. 2100472. Available at: 10.1002/smll.202100472.

Gowers, R.J. et al. (2016) ‘MDAnalysis: A Python Package for the Rapid Analysis of Molecular Dynamics Simulations’, Proceedings of the 15th Python in Science Conference, pp. 98–105. Available at: 10.25080/Majora-629e541a-00e.

Guntas, G. et al. (2005) ‘Directed evolution of protein switches and their application to the creation of ligand-binding proteins’, Proceedings of the National Academy of Sciences, 102(32), pp. 11224–11229. Available at: 10.1073/pnas.0502673102.

Guntas, G. and Ostermeier, M. (2004) ‘Creation of an Allosteric Enzyme by Domain Insertion’, Journal of Molecular Biology, 336(1), pp. 263–273. Available at: 10.1016/j.jmb.2003.12.016.

Hu, H. et al. (2018) ‘Glucose monitoring in living cells with single fluorescent protein-based sensors’, RSC Advances, 8(5), pp. 2485–2489. Available at: 10.1039/C7RA11347A.

Jing, M. et al. (2020) ‘An optimized acetylcholine sensor for monitoring in vivo cholinergic activity’, Nature Methods, 17(11), pp. 1139–1146. Available at: 10.1038/s41592-020-0953-2.

Li, H. et al. (2022) ‘Tailoring Escherichia coli Chemotactic Sensing towards Cadmium by Computational Redesign of Ribose-Binding Protein’, mSystems, 7(1), pp. e01084–21. Available at: 10.1128/msystems.01084-21.

Marvin, J.S. et al. (2011) ‘A genetically encoded, high-signal-to-noise maltose sensor’, Proteins: Structure, Function, and Bioinformatics, 79(11), pp. 3025–3036. Available at: 10.1002/prot.23118.

Marvin, J.S. et al. (2013) ‘An optimized fluorescent probe for visualizing glutamate neurotransmission’, Nature Methods, 10(2), pp. 162–170. Available at: 10.1038/nmeth.2333.

Marvin, J.S. et al. (2019) ‘A genetically encoded fluorescent sensor for in vivo imaging of GABA’, Nature Methods, 16(8), pp. 763–770. Available at: 10.1038/s41592-019-0471-2.

McGibbon, R.T. et al. (2015) ‘MDTraj: A Modern Open Library for the Analysis of Molecular Dynamics Trajectories’, Biophysical Journal, 109(8), pp. 1528–1532. Available at: 10.1016/j.bpj.2015.08.015.

Mercadante, D., Gräter, F. and Daday, C. (2018) ‘CONAN: A Tool to Decode Dynamical Information from Molecular Interaction Maps’, Biophysical Journal, 114(6), pp. 1267–1273. Available at: 10.1016/j.bpj.2018.01.033.

Messerli, M.A. and Robinson, K.R. (1998) ‘Cytoplasmic acidification and current influx follow growth pulses of Lilium longiflorum pollen tubes’, The Plant Journal, 16(1), pp. 87–91. Available at: 10.1046/j.1365-313x.1998.00266.x.

Mukherjee, P. et al. (2015) ‘Live imaging of inorganic phosphate in plants with cellular and subcellular resolution’, Plant Physiology, 167(3), pp. 628–638. Available at: 10.1104/pp.114.254003.

Nadler, D.C. et al. (2016) ‘Rapid construction of metabolite biosensors using domain-insertion profiling’, Nature Communications, 7(1), p. 12266. Available at: 10.1038/ncomms12266.

Nasu, Y. et al. (2021) ‘Structure- and mechanism-guided design of single fluorescent protein-based biosensors’, Nature chemical biology, 17(5), pp. 509–518. Available at: 10.1038/s41589-020-00718-x.

Praetorius, F. et al. (2023) ‘Design of stimulus-responsive two-state hinge proteins’, Science, 381(6659), pp. 754–760. Available at: 10.1126/science.adg7731.

Quiocho, F.A. and Ledvina, P.S. (1996) ‘Atomic structure and specificity of bacterial periplasmic receptors for active transport and chemotaxis: variation of common themes’, Molecular Microbiology, 20(1), pp. 17–25. Available at: 10.1111/j.1365-2958.1996.tb02484.x.

Quiocho, F.A., Spurlino, J.C. and Rodseth, L.E. (1997) ‘Extensive features of tight oligosaccharide binding revealed in high-resolution structures of the maltodextrin transport/chemosensory receptor’, Structure (London, England: 1993), 5(8), pp. 997–1015. Available at: 10.1016/s0969-2126(97)00253-0.

Ribeiro, L.F. et al. (2015) ‘Insertion of a xylanase in xylose binding protein results in a xylose-stimulated xylanase’, Biotechnology for Biofuels, 8(1), p. 118. Available at: 10.1186/s13068-015-0293-0.

Ribeiro, L.F. et al. (2016) ‘A xylose-stimulated xylanase–xylose binding protein chimera created by random nonhomologous recombination’, Biotechnology for Biofuels, 9(1), p. 119. Available at: 10.1186/s13068-016-0529-7.

Ribeiro, L.F. et al. (2019) ‘Converting a Periplasmic Binding Protein into a Synthetic Biosensing Switch through Domain Insertion’, BioMed Research International, 2019, p. 4798793. Available at: 10.1155/2019/4798793.

Saldaño, T. et al./person-group>. (2022) ‘Impact of protein conformational diversity on AlphaFold predictions’, Bioinformatics. Edited by A. Valencia, 38(10), pp. 2742–2748. Available at: 10.1093/bioinformatics/btac202.

Shivange, A.V. et al. (2019) ‘Determining the pharmacokinetics of nicotinic drugs in the endoplasmic reticulum using biosensors’, The Journal of General Physiology, 151(6), pp. 738–757. Available at: 10.1085/jgp.201812201.

Tavares, D. et al. (2019) ‘Computational redesign of the Escherichia coli ribose-binding protein ligand binding pocket for 1,3-cyclohexanediol and cyclohexanol’, Scientific Reports, 9(1), p. 16940. Available at: 10.1038/s41598-019-53507-5.

Taylor, N.D. et al. (2016) ‘Engineering an allosteric transcription factor to respond to new ligands’, Nature Methods, 13(2), pp. 177–183. Available at: 10.1038/nmeth.3696.

Younger, A.K.D. et al. (2017) ‘Engineering Modular Biosensors to Confer Metabolite-Responsive Regulation of Transcription’, ACS Synthetic Biology, 6(2), pp. 311–325. Available at: 10.1021/acssynbio.6b00184.

Younger, A.K.D. et al. (2018) ‘Development of novel metabolite-responsive transcription factors via transposon-mediated protein fusion’, Protein Engineering, Design and Selection, 31(2), pp. 55–63. Available at: 10.1093/protein/gzy001.

Yu, Y. and Lutz, S. (2011) ‘Circular permutation: a different way to engineer enzyme structure and function’, Trends in Biotechnology, 29(1), pp. 18–25. Available at: 10.1016/j.tibtech.2010.10.004.

