## Supplementary Material for "Computational Design of Periplasmic Binding Protein Biosensors Guided by Molecular Dynamics"

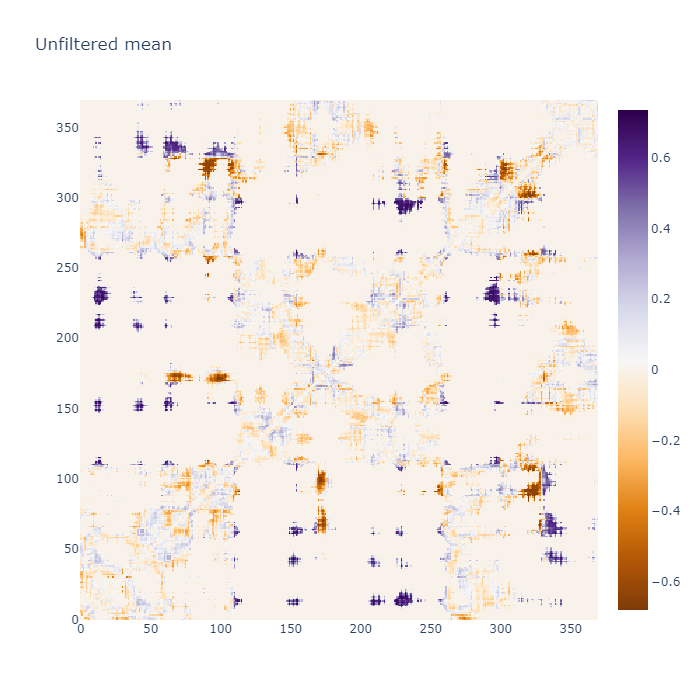

**Supplementary figure 1.** Unfiltered matrix of mean Pearson Correlation Coefficient values for inter-residue contacts over all apo simulations where the closed-to-open transition occurred.

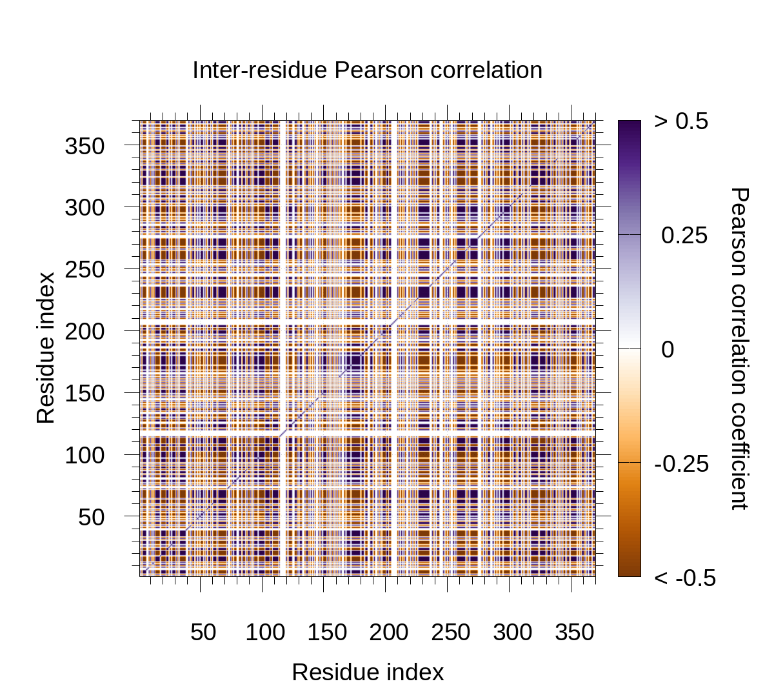

**Supplementary figure 2.** Pearson Correlation Coefficient matrix of inter-residue contacts calculated from comparison of the available crystal structures of apo (1ANF) and holo (1OMP) MBP.

|  | **Raw fluorescence** | | | | | | | | | | | |
| --- | --- | --- | --- | --- | --- | --- | --- | --- | --- | --- | --- | --- |
|  | **0 mM** | | | | **1 mM** | | | | **100 mM** | | | |
| **Sensor** | **replicate 1** | **replicate 2** | **replicate 3** | **mean** | **replicate 1** | **replicate 2** | **replicate 3** | **mean** | **replicate 1** | **replicate 2** | **replicate 3** | **mean** |
| cpEGFP | 609 | 1818 | 1932 | 1453 | 637 | 1781 | 1834 | 1417 | 783 | 1753 | 1851 | 1462 |
| MBP | -1 | -2 | -2 | -2 | -1 | -2 | -3 | -2 | -1 | -2 | -3 | -2 |
| 170 | 460 | 477 | 521 | 486 | 689 | 687 | 807 | 727 | 707 | 700 | 818 | 742 |
| 348 | 1667 | 1490 | 1107 | 1421 | 2257 | 2211 | 1656 | 2041 | 2836 | 2387 | 1823 | 2349 |
| 356 | 110 | 179 | 126 | 138 | 167 | 190 | 146 | 168 | 192 | 198 | 171 | 187 |
| 205 | 320 | 324 | 349 | 331 | 308 | 259 | 277 | 281 | 286 | 275 | 341 | 300 |
| 208 | 460 | 456 | 455 | 457 | 484 | 379 | 435 | 432 | 400 | 446 | 428 | 425 |
| 211 | 114 | 102 | 116 | 111 | 110 | 93 | 105 | 103 | 103 | 93 | 119 | 105 |
| 214 | 297 | 276 | 252 | 275 | 240 | 177 | 218 | 212 | 303 | 204 | 201 | 236 |
| 217 | 433 | 423 | 465 | 440 | 637 | 443 | 603 | 561 | 549 | 614 | 648 | 604 |
| 224 | 7 | 3 | 9 | 6 | 7 | 4 | 11 | 7 | 6 | 9 | 11 | 9 |
| 227 | 15 | 13 | 20 | 16 | 19 | 14 | 21 | 18 | 18 | 21 | 23 | 21 |
| 230 | 17 | 15 | 22 | 18 | 24 | 21 | 31 | 25 | 72 | 77 | 120 | 90 |
| 233 | 134 | 113 | 130 | 126 | 260 | 186 | 210 | 218 | 214 | 204 | 238 | 219 |
| 236 | 255 | 190 | 229 | 225 | 283 | 199 | 241 | 241 | 284 | 211 | 295 | 263 |
| 131 | 227 | 282 | 264 | 258 | 222 | 237 | 225 | 228 | 213 | 201 | 179 | 197 |
| 195 | 327 | 447 | 350 | 375 | 334 | 388 | 301 | 341 | 327 | 276 | 241 | 281 |

**Supplementary table 1.** Mean raw fluorescence values for all sensors and controls. “Replicate” refers to “biological replicate”, each being a mean of three technical repeats (technical repeat values not shown).

|  | **dF/F0** | | | | | | | |
| --- | --- | --- | --- | --- | --- | --- | --- | --- |
|  | **1 mM** | | | | **100 mM** | | | |
| **Sensor** | **replicate 1** | **replicate 2** | **replicate 3** | **mean** | **replicate 1** | **replicate 2** | **replicate 3** | **mean** |
| cpEGFP | 0.047 | -0.02 | 0.262 | 0.096 | 0.286 | -0.036 | 0.274 | 0.175 |
| MBP | 0 | 0 | 0.6 | 0.2 | 1 | 0 | 0.8 | 0.6 |
| 170 | 0.498 | 0.439 | 0.547 | 0.495 | 0.537 | 0.467 | 0.57 | 0.525 |
| 348 | 0.354 | 0.484 | 0.495 | 0.445 | 0.702 | 0.602 | 0.646 | 0.65 |
| 356 | 0.523 | 0.061 | 0.161 | 0.249 | 0.748 | 0.106 | 0.36 | 0.405 |
| 205 | -0.039 | -0.201 | -0.206 | -0.149 | -0.108 | -0.153 | -0.024 | -0.095 |
| 208 | 0.051 | -0.169 | -0.044 | -0.054 | -0.13 | -0.021 | -0.058 | -0.07 |
| 211 | -0.041 | -0.085 | -0.089 | -0.072 | -0.102 | -0.088 | 0.026 | -0.055 |
| 214 | -0.193 | -0.358 | -0.135 | -0.229 | 0.02 | -0.26 | -0.201 | -0.147 |
| 217 | 0.47 | 0.047 | 0.296 | 0.271 | 0.268 | 0.452 | 0.394 | 0.371 |
| 224 | 0 | 0.5 | 0.214 | 0.238 | -0.095 | 2.25 | 0.143 | 0.766 |
| 227 | 0.217 | 0.132 | 0.033 | 0.127 | 0.196 | 0.632 | 0.148 | 0.325 |
| 230 | 0.365 | 0.378 | 0.431 | 0.391 | 3.154 | 4.156 | 4.554 | 3.954 |
| 233 | 0.935 | 0.638 | 0.612 | 0.729 | 0.591 | 0.803 | 0.828 | 0.741 |
| 236 | 0.108 | 0.046 | 0.054 | 0.069 | 0.112 | 0.109 | 0.289 | 0.17 |
| 131 | -0.025 | -0.16 | -0.147 | -0.11 | -0.065 | -0.288 | -0.322 | -0.225 |
| 195 | 0.021 | -0.131 | -0.141 | -0.083 | -0.002 | -0.381 | -0.31 | -0.231 |

**Supplementary table 2.** dF/F0 values for all sensors and controls.

**Supplementary table 3 (below).** FASTA format nucleotide and amino acid sequences used in this study. Maltose binding protein sequence (MBP) is highlighted in blue, circularly permuted GFP sequence (cpGFP) in yellow, and linker sequence between these two components in biosensors in green.

>MBP nucleotide

ATGAAAATCGAAGAAGGTAAACTGGTAATCTGGATTAACGGCGATAAAGGCTATAACGGACTCGCTGAA

GTCGGTAAGAAATTCGAGAAAGATACCGGAATTAAAGTCACCGTTGAGCATCCGGATAAACTGGAAGAG

AAATTCCCACAGGTTGCGGCAACTGGCGATGGCCCTGACATTATCTTCTGGGCACACGACCGCTTTGGT

GGCTACGCTCAATCTGGCCTGTTGGCTGAAATCACCCCGGACAAAGCGTTCCAGGACAAGCTGTATCCG

TTTACCTGGGATGCCGTACGTTACAACGGCAAGCTGATTGCTTACCCGATCGCTGTTGAAGCGTTATCG

CTGATTTATAACAAAGATCTGCTGCCGAACCCGCCAAAAACCTGGGAAGAGATCCCGGCGCTGGATAAA

GAACTGAAAGCGAAAGGTAAGAGCGCGCTGATGTTCAACCTGCAAGAACCGTACTTCACCTGGCCGCTG

ATTGCTGCTGACGGGGGTTATGCGTTCAAGTATGAAAACGGCAAGTACGACATTAAAGACGTGGGCGTG

GATAACGCTGGCGCGAAAGCGGGTCTGACCTTCCTGGTTGACCTGATTAAAAACAAACACATGAATGCA

GACACCGATTACTCCATCGCAGAAGCTGCCTTTAATAAAGGCGAAACAGCGATGACCATCAACGGCCCG

TGGGCATGGTCCAACATCGACACCAGCAAAGTGAATTATGGTGTAACGGTACTGCCGACCTTCAAGGGT

CAACCATCCAAACCGTTCGTTGGCGTGCTGAGCGCAGGTATTAACGCCGCCAGTCCGAACAAAGAGCTG

GCGAAAGAGTTCCTCGAAAACTATCTGCTGACTGATGAAGGTCTGGAAGCGGTTAATAAAGACAAACCG

CTGGGTGCCGTAGCGCTGAAGTCTTACGAGGAAGAGTTGGCGAAAGATCCACGTATTGCCGCCACCATG

GAAAACGCCCAGAAAGGTGAAATCATGCCGAACATCCCGCAGATGTCCGCTTTCTGGTATGCCGTGCGT

ACTGCGGTGATCAACGCCGCCAGCGGTCGTCAGACTGTCGATGAAGCCCTGAAAGACGCGCAGACTCGT

ATCACCAAGTAA

>MBP amino acid

MKIEEGKLVIWINGDKGYNGLAEVGKKFEKDTGIKVTVEHPDKLEEKFPQVAATGDGPDIIFWAHDRFG

GYAQSGLLAEITPDKAFQDKLYPFTWDAVRYNGKLIAYPIAVEALSLIYNKDLLPNPPKTWEEIPALDK

ELKAKGKSALMFNLQEPYFTWPLIAADGGYAFKYENGKYDIKDVGVDNAGAKAGLTFLVDLIKNKHMNA

DTDYSIAEAAFNKGETAMTINGPWAWSNIDTSKVNYGVTVLPTFKGQPSKPFVGVLSAGINAASPNKEL

AKEFLENYLLTDEGLEAVNKDKPLGAVALKSYEEELAKDPRIAATMENAQKGEIMPNIPQMSAFWYAVR

TAVINAASGRQTVDEALKDAQTRITK*

>cpGFP nucleotide

ATGTATAACGTCTTTATCATGGCCGACAAGCAGAAGAACGGCATCAAGGCGAACTTCAAGATCCGCCAC

AACATCGAGGACGGCGGCGTGCAGCTCGCCTATCACTACCAGCAGAACACCCCCATCGGCGACGGCCCC

GTGCTGCTGCCCGACAACCACTACCTGAGCGTGCAGTCCAAACTGAGCAAAGACCCCAACGAGAAGCGC

GATCACATGGTCCTGCTGGAGTTCGTGACCGCCGCCGGGATCACTCTCGGCATGGACGAGCTGTACAAG

GGCGGTACCGGAGGGAGCATGGTGAGCAAGGGCGAGGAGCTGTTCACCGGGGTGGTGCCCATCCTGGTC

GAGCTGGACGGCGACGTAAACGGCCACAAGTTCAGCGTGTCCGGCGAGGGCGAGGGCGATGCCACCTAC

GGCAAGCTGACCCTGAAGTTCATCTGCACCACCGGCAAGCTGCCCGTGCCCTGGCCCACCCTCGTGACC

ACCCTGACCTACGGCGTGCAGTGCTTCAGCCGCTACCCCGACCACATGAAGCAGCACGACTTCTTCAAG

TCCGCCATGCCCGAAGGCTACATTCAGGAGCGCACCATCTTCTTCAAGGACGACGGCAACTATAAGACA

CGCGCTGAGGTTAAGTTCGAGGGCGACACTCTGGTTAACCGCATCGAGCTGAAGGGCATCGACTTCAAG

GAGGACGGCAACATCCTGGGCCATAAGCTTGAATATAACTTCAACTAA

>cpGFP amino acid

MYNVFIMADKQKNGIKANFKIRHNIEDGGVQLAYHYQQNTPIGDGPVLLPDNHYLSVQSKLSKDPNEKR

DHMVLLEFVTAAGITLGMDELYKGGTGGSMVSKGEELFTGVVPILVELDGDVNGHKFSVSGEGEGDATY

GKLTLKFICTTGKLPVPWPTLVTTLTYGVQCFSRYPDHMKQHDFFKSAMPEGYIQERTIFFKDDGNYKT

RAEVKFEGDTLVNRIELKGIDFKEDGNILGHKLEYNFN*

>MBP_cpEGFP170 nucleotide

ATGAAAATCGAAGAAGGTAAACTGGTAATCTGGATTAACGGCGATAAAGGCTATAACGGACTCGCTGAA

GTCGGTAAGAAATTCGAGAAAGATACCGGAATTAAAGTCACCGTTGAGCATCCGGATAAACTGGAAGAG

AAATTCCCACAGGTTGCGGCAACTGGCGATGGCCCTGACATTATCTTCTGGGCACACGACCGCTTTGGT

GGCTACGCTCAATCTGGCCTGTTGGCTGAAATCACCCCGGACAAAGCGTTCCAGGACAAGCTGTATCCG

TTTACCTGGGATGCCGTACGTTACAACGGCAAGCTGATTGCTTACCCGATCGCTGTTGAAGCGTTATCG

CTGATTTATAACAAAGATCTGCTGCCGAACCCGCCAAAAACCTGGGAAGAGATCCCGGCGCTGGATAAA

GAACTGAAAGCGAAAGGTAAGAGCGCGCTGATGTTCAACCTGCAAGAACCGTACTTCACCTGGCCGCTG

ATTGCTGCTGACGGGGGTTATGCGTTCAAGTATGCATCTTATAACGTCTTTATCATGGCCGACAAGCAG

AAGAACGGCATCAAGGCGAACTTCAAGATCCGCCACAACATCGAGGACGGCGGCGTGCAGCTCGCCTAT

CACTACCAGCAGAACACCCCCATCGGCGACGGCCCCGTGCTGCTGCCCGACAACCACTACCTGAGCGTG

CAGTCCAAACTGAGCAAAGACCCCAACGAGAAGCGCGATCACATGGTCCTGCTGGAGTTCGTGACCGCC

GCCGGGATCACTCTCGGCATGGACGAGCTGTACAAGGGCGGTACCGGAGGGAGCATGGTGAGCAAGGGC

GAGGAGCTGTTCACCGGGGTGGTGCCCATCCTGGTCGAGCTGGACGGCGACGTAAACGGCCACAAGTTC

AGCGTGTCCGGCGAGGGCGAGGGCGATGCCACCTACGGCAAGCTGACCCTGAAGTTCATCTGCACCACC

GGCAAGCTGCCCGTGCCCTGGCCCACCCTCGTGACCACCCTGACCTACGGCGTGCAGTGCTTCAGCCGC

TACCCCGACCACATGAAGCAGCACGACTTCTTCAAGTCCGCCATGCCCGAAGGCTACATTCAGGAGCGC

ACCATCTTCTTCAAGGACGACGGCAACTATAAGACACGCGCTGAGGTTAAGTTCGAGGGCGACACTCTG

GTTAACCGCATCGAGCTGAAGGGCATCGACTTCAAGGAGGACGGCAACATCCTGGGCCATAAGCTTGAA

TATAACTTCAACGCATCTAAGTATGAAAACGGCAAGTACGACATTAAAGACGTGGGCGTGGATAACGCT

GGCGCGAAAGCGGGTCTGACCTTCCTGGTTGACCTGATTAAAAACAAACACATGAATGCAGACACCGAT

TACTCCATCGCAGAAGCTGCCTTTAATAAAGGCGAAACAGCGATGACCATCAACGGCCCGTGGGCATGG

TCCAACATCGACACCAGCAAAGTGAATTATGGTGTAACGGTACTGCCGACCTTCAAGGGTCAACCATCC

AAACCGTTCGTTGGCGTGCTGAGCGCAGGTATTAACGCCGCCAGTCCGAACAAAGAGCTGGCGAAAGAG

TTCCTCGAAAACTATCTGCTGACTGATGAAGGTCTGGAAGCGGTTAATAAAGACAAACCGCTGGGTGCC

GTAGCGCTGAAGTCTTACGAGGAAGAGTTGGCGAAAGATCCACGTATTGCCGCCACCATGGAAAACGCC

CAGAAAGGTGAAATCATGCCGAACATCCCGCAGATGTCCGCTTTCTGGTATGCCGTGCGTACTGCGGTG

ATCAACGCCGCCAGCGGTCGTCAGACTGTCGATGAAGCCCTGAAAGACGCGCAGACTCGTATCACCAAG

TAA

>MBP_cpEGFP170 amino acid

MKIEEGKLVIWINGDKGYNGLAEVGKKFEKDTGIKVTVEHPDKLEEKFPQVAATGDGPDIIFWAHDRFG

GYAQSGLLAEITPDKAFQDKLYPFTWDAVRYNGKLIAYPIAVEALSLIYNKDLLPNPPKTWEEIPALDK

ELKAKGKSALMFNLQEPYFTWPLIAADGGYAFKYASYNVFIMADKQKNGIKANFKIRHNIEDGGVQLAY

HYQQNTPIGDGPVLLPDNHYLSVQSKLSKDPNEKRDHMVLLEFVTAAGITLGMDELYKGGTGGSMVSKG

EELFTGVVPILVELDGDVNGHKFSVSGEGEGDATYGKLTLKFICTTGKLPVPWPTLVTTLTYGVQCFSR

YPDHMKQHDFFKSAMPEGYIQERTIFFKDDGNYKTRAEVKFEGDTLVNRIELKGIDFKEDGNILGHKLE

YNFNASKYENGKYDIKDVGVDNAGAKAGLTFLVDLIKNKHMNADTDYSIAEAAFNKGETAMTINGPWAW

SNIDTSKVNYGVTVLPTFKGQPSKPFVGVLSAGINAASPNKELAKEFLENYLLTDEGLEAVNKDKPLGA

VALKSYEEELAKDPRIAATMENAQKGEIMPNIPQMSAFWYAVRTAVINAASGRQTVDEALKDAQTRITK

*

>MBP_cpEGFP348 nucleotide ATGAAAATCGAAGAAGGTAAACTGGTAATCTGGATTAACGGCGATAAAGGCTATAACGGACTCGCTGAA

GTCGGTAAGAAATTCGAGAAAGATACCGGAATTAAAGTCACCGTTGAGCATCCGGATAAACTGGAAGAG

AAATTCCCACAGGTTGCGGCAACTGGCGATGGCCCTGACATTATCTTCTGGGCACACGACCGCTTTGGT

GGCTACGCTCAATCTGGCCTGTTGGCTGAAATCACCCCGGACAAAGCGTTCCAGGACAAGCTGTATCCG

TTTACCTGGGATGCCGTACGTTACAACGGCAAGCTGATTGCTTACCCGATCGCTGTTGAAGCGTTATCG

CTGATTTATAACAAAGATCTGCTGCCGAACCCGCCAAAAACCTGGGAAGAGATCCCGGCGCTGGATAAA

GAACTGAAAGCGAAAGGTAAGAGCGCGCTGATGTTCAACCTGCAAGAACCGTACTTCACCTGGCCGCTG

ATTGCTGCTGACGGGGGTTATGCGTTCAAGTATGAAAACGGCAAGTACGACATTAAAGACGTGGGCGTG

GATAACGCTGGCGCGAAAGCGGGTCTGACCTTCCTGGTTGACCTGATTAAAAACAAACACATGAATGCA

GACACCGATTACTCCATCGCAGAAGCTGCCTTTAATAAAGGCGAAACAGCGATGACCATCAACGGCCCG

TGGGCATGGTCCAACATCGACACCAGCAAAGTGAATTATGGTGTAACGGTACTGCCGACCTTCAAGGGT

CAACCATCCAAACCGTTCGTTGGCGTGCTGAGCGCAGGTATTAACGCCGCCAGTCCGAACAAAGAGCTG

GCGAAAGAGTTCCTCGAAAACTATCTGCTGACTGATGAAGGTCTGGAAGCGGTTAATAAAGACAAACCG

CTGGGTGCCGTAGCGCTGAAGTCTTACGAGGAAGAGTTGGCGAAAGATCCACGTATTGCCGCCACCATG

GAAAACGCCCAGAAAGGTGAAATCATGCCGAACATCCCGCAGATGTCCGCTTTCTGGTATGCCGTGCGT

ACTGCGGTGATCAACGCATCTTATAACGTCTTTATCATGGCCGACAAGCAGAAGAACGGCATCAAGGCG

AACTTCAAGATCCGCCACAACATCGAGGACGGCGGCGTGCAGCTCGCCTATCACTACCAGCAGAACACC

CCCATCGGCGACGGCCCCGTGCTGCTGCCCGACAACCACTACCTGAGCGTGCAGTCCAAACTGAGCAAA

GACCCCAACGAGAAGCGCGATCACATGGTCCTGCTGGAGTTCGTGACCGCCGCCGGGATCACTCTCGGC

ATGGACGAGCTGTACAAGGGCGGTACCGGAGGGAGCATGGTGAGCAAGGGCGAGGAGCTGTTCACCGGG

GTGGTGCCCATCCTGGTCGAGCTGGACGGCGACGTAAACGGCCACAAGTTCAGCGTGTCCGGCGAGGGC

GAGGGCGATGCCACCTACGGCAAGCTGACCCTGAAGTTCATCTGCACCACCGGCAAGCTGCCCGTGCCC

TGGCCCACCCTCGTGACCACCCTGACCTACGGCGTGCAGTGCTTCAGCCGCTACCCCGACCACATGAAG

CAGCACGACTTCTTCAAGTCCGCCATGCCCGAAGGCTACATTCAGGAGCGCACCATCTTCTTCAAGGAC

GACGGCAACTATAAGACACGCGCTGAGGTTAAGTTCGAGGGCGACACTCTGGTTAACCGCATCGAGCTG

AAGGGCATCGACTTCAAGGAGGACGGCAACATCCTGGGCCATAAGCTTGAATATAACTTCAACGCATCT

ATCAACGCCGCCAGCGGTCGTCAGACTGTCGATGAAGCCCTGAAAGACGCGCAGACTCGTATCACCAAG

TAA

>MBP_cpEGFP348 amino acid

MKIEEGKLVIWINGDKGYNGLAEVGKKFEKDTGIKVTVEHPDKLEEKFPQVAATGDGPDIIFWAHDRFG

GYAQSGLLAEITPDKAFQDKLYPFTWDAVRYNGKLIAYPIAVEALSLIYNKDLLPNPPKTWEEIPALDK

ELKAKGKSALMFNLQEPYFTWPLIAADGGYAFKYENGKYDIKDVGVDNAGAKAGLTFLVDLIKNKHMNA

DTDYSIAEAAFNKGETAMTINGPWAWSNIDTSKVNYGVTVLPTFKGQPSKPFVGVLSAGINAASPNKEL

AKEFLENYLLTDEGLEAVNKDKPLGAVALKSYEEELAKDPRIAATMENAQKGEIMPNIPQMSAFWYAVR

TAVINASYNVFIMADKQKNGIKANFKIRHNIEDGGVQLAYHYQQNTPIGDGPVLLPDNHYLSVQSKLSK

DPNEKRDHMVLLEFVTAAGITLGMDELYKGGTGGSMVSKGEELFTGVVPILVELDGDVNGHKFSVSGEG

EGDATYGKLTLKFICTTGKLPVPWPTLVTTLTYGVQCFSRYPDHMKQHDFFKSAMPEGYIQERTIFFKD

DGNYKTRAEVKFEGDTLVNRIELKGIDFKEDGNILGHKLEYNFNASINAASGRQTVDEALKDAQTRITK

*

>MBP_cpEGFP205 nucleotide ATGAAAATCGAAGAAGGTAAACTGGTAATCTGGATTAACGGCGATAAAGGCTATAACGGACTCGCTGAA

GTCGGTAAGAAATTCGAGAAAGATACCGGAATTAAAGTCACCGTTGAGCATCCGGATAAACTGGAAGAG

AAATTCCCACAGGTTGCGGCAACTGGCGATGGCCCTGACATTATCTTCTGGGCACACGACCGCTTTGGT

GGCTACGCTCAATCTGGCCTGTTGGCTGAAATCACCCCGGACAAAGCGTTCCAGGACAAGCTGTATCCG

TTTACCTGGGATGCCGTACGTTACAACGGCAAGCTGATTGCTTACCCGATCGCTGTTGAAGCGTTATCG

CTGATTTATAACAAAGATCTGCTGCCGAACCCGCCAAAAACCTGGGAAGAGATCCCGGCGCTGGATAAA

GAACTGAAAGCGAAAGGTAAGAGCGCGCTGATGTTCAACCTGCAAGAACCGTACTTCACCTGGCCGCTG

ATTGCTGCTGACGGGGGTTATGCGTTCAAGTATGAAAACGGCAAGTACGACATTAAAGACGTGGGCGTG

GATAACGCTGGCGCGAAAGCGGGTCTGACCTTCCTGGTTGACCTGATTAAAAACAAACACATGAATGCA

GCATCTTATAACGTCTTTATCATGGCCGACAAGCAGAAGAACGGCATCAAGGCGAACTTCAAGATCCGC

CACAACATCGAGGACGGCGGCGTGCAGCTCGCCTATCACTACCAGCAGAACACCCCCATCGGCGACGGC

CCCGTGCTGCTGCCCGACAACCACTACCTGAGCGTGCAGTCCAAACTGAGCAAAGACCCCAACGAGAAG

CGCGATCACATGGTCCTGCTGGAGTTCGTGACCGCCGCCGGGATCACTCTCGGCATGGACGAGCTGTAC

AAGGGCGGTACCGGAGGGAGCATGGTGAGCAAGGGCGAGGAGCTGTTCACCGGGGTGGTGCCCATCCTG

GTCGAGCTGGACGGCGACGTAAACGGCCACAAGTTCAGCGTGTCCGGCGAGGGCGAGGGCGATGCCACC

TACGGCAAGCTGACCCTGAAGTTCATCTGCACCACCGGCAAGCTGCCCGTGCCCTGGCCCACCCTCGTG

ACCACCCTGACCTACGGCGTGCAGTGCTTCAGCCGCTACCCCGACCACATGAAGCAGCACGACTTCTTC

AAGTCCGCCATGCCCGAAGGCTACATTCAGGAGCGCACCATCTTCTTCAAGGACGACGGCAACTATAAG

ACACGCGCTGAGGTTAAGTTCGAGGGCGACACTCTGGTTAACCGCATCGAGCTGAAGGGCATCGACTTC

AAGGAGGACGGCAACATCCTGGGCCATAAGCTTGAATATAACTTCAACGCATCTAATGCAGACACCGAT

TACTCCATCGCAGAAGCTGCCTTTAATAAAGGCGAAACAGCGATGACCATCAACGGCCCGTGGGCATGG

TCCAACATCGACACCAGCAAAGTGAATTATGGTGTAACGGTACTGCCGACCTTCAAGGGTCAACCATCC

AAACCGTTCGTTGGCGTGCTGAGCGCAGGTATTAACGCCGCCAGTCCGAACAAAGAGCTGGCGAAAGAG

TTCCTCGAAAACTATCTGCTGACTGATGAAGGTCTGGAAGCGGTTAATAAAGACAAACCGCTGGGTGCC

GTAGCGCTGAAGTCTTACGAGGAAGAGTTGGCGAAAGATCCACGTATTGCCGCCACCATGGAAAACGCC

CAGAAAGGTGAAATCATGCCGAACATCCCGCAGATGTCCGCTTTCTGGTATGCCGTGCGTACTGCGGTG

ATCAACGCCGCCAGCGGTCGTCAGACTGTCGATGAAGCCCTGAAAGACGCGCAGACTCGTATCACCAAG

TAA

>MBP_cpEGFP205 amino acid

MKIEEGKLVIWINGDKGYNGLAEVGKKFEKDTGIKVTVEHPDKLEEKFPQVAATGDGPDIIFWAHDRFG

GYAQSGLLAEITPDKAFQDKLYPFTWDAVRYNGKLIAYPIAVEALSLIYNKDLLPNPPKTWEEIPALDK

ELKAKGKSALMFNLQEPYFTWPLIAADGGYAFKYENGKYDIKDVGVDNAGAKAGLTFLVDLIKNKHMNA

ASYNVFIMADKQKNGIKANFKIRHNIEDGGVQLAYHYQQNTPIGDGPVLLPDNHYLSVQSKLSKDPNEK

RDHMVLLEFVTAAGITLGMDELYKGGTGGSMVSKGEELFTGVVPILVELDGDVNGHKFSVSGEGEGDAT

YGKLTLKFICTTGKLPVPWPTLVTTLTYGVQCFSRYPDHMKQHDFFKSAMPEGYIQERTIFFKDDGNYK

TRAEVKFEGDTLVNRIELKGIDFKEDGNILGHKLEYNFNASNADTDYSIAEAAFNKGETAMTINGPWAW

SNIDTSKVNYGVTVLPTFKGQPSKPFVGVLSAGINAASPNKELAKEFLENYLLTDEGLEAVNKDKPLGA

VALKSYEEELAKDPRIAATMENAQKGEIMPNIPQMSAFWYAVRTAVINAASGRQTVDEALKDAQTRITK

*

>MBP_cpEGFP208 nucleotide

ATGAAAATCGAAGAAGGTAAACTGGTAATCTGGATTAACGGCGATAAAGGCTATAACGGACTCGCTGAA

GTCGGTAAGAAATTCGAGAAAGATACCGGAATTAAAGTCACCGTTGAGCATCCGGATAAACTGGAAGAG

AAATTCCCACAGGTTGCGGCAACTGGCGATGGCCCTGACATTATCTTCTGGGCACACGACCGCTTTGGT

GGCTACGCTCAATCTGGCCTGTTGGCTGAAATCACCCCGGACAAAGCGTTCCAGGACAAGCTGTATCCG

TTTACCTGGGATGCCGTACGTTACAACGGCAAGCTGATTGCTTACCCGATCGCTGTTGAAGCGTTATCG

CTGATTTATAACAAAGATCTGCTGCCGAACCCGCCAAAAACCTGGGAAGAGATCCCGGCGCTGGATAAA

GAACTGAAAGCGAAAGGTAAGAGCGCGCTGATGTTCAACCTGCAAGAACCGTACTTCACCTGGCCGCTG

ATTGCTGCTGACGGGGGTTATGCGTTCAAGTATGAAAACGGCAAGTACGACATTAAAGACGTGGGCGTG

GATAACGCTGGCGCGAAAGCGGGTCTGACCTTCCTGGTTGACCTGATTAAAAACAAACACATGAATGCA

GACACCGATGCATCTTATAACGTCTTTATCATGGCCGACAAGCAGAAGAACGGCATCAAGGCGAACTTC

AAGATCCGCCACAACATCGAGGACGGCGGCGTGCAGCTCGCCTATCACTACCAGCAGAACACCCCCATC

GGCGACGGCCCCGTGCTGCTGCCCGACAACCACTACCTGAGCGTGCAGTCCAAACTGAGCAAAGACCCC

AACGAGAAGCGCGATCACATGGTCCTGCTGGAGTTCGTGACCGCCGCCGGGATCACTCTCGGCATGGAC

GAGCTGTACAAGGGCGGTACCGGAGGGAGCATGGTGAGCAAGGGCGAGGAGCTGTTCACCGGGGTGGTG

CCCATCCTGGTCGAGCTGGACGGCGACGTAAACGGCCACAAGTTCAGCGTGTCCGGCGAGGGCGAGGGC

GATGCCACCTACGGCAAGCTGACCCTGAAGTTCATCTGCACCACCGGCAAGCTGCCCGTGCCCTGGCCC

ACCCTCGTGACCACCCTGACCTACGGCGTGCAGTGCTTCAGCCGCTACCCCGACCACATGAAGCAGCAC

GACTTCTTCAAGTCCGCCATGCCCGAAGGCTACATTCAGGAGCGCACCATCTTCTTCAAGGACGACGGC

AACTATAAGACACGCGCTGAGGTTAAGTTCGAGGGCGACACTCTGGTTAACCGCATCGAGCTGAAGGGC

ATCGACTTCAAGGAGGACGGCAACATCCTGGGCCATAAGCTTGAATATAACTTCAACGCATCTACCGAT

TACTCCATCGCAGAAGCTGCCTTTAATAAAGGCGAAACAGCGATGACCATCAACGGCCCGTGGGCATGG

TCCAACATCGACACCAGCAAAGTGAATTATGGTGTAACGGTACTGCCGACCTTCAAGGGTCAACCATCC

AAACCGTTCGTTGGCGTGCTGAGCGCAGGTATTAACGCCGCCAGTCCGAACAAAGAGCTGGCGAAAGAG

TTCCTCGAAAACTATCTGCTGACTGATGAAGGTCTGGAAGCGGTTAATAAAGACAAACCGCTGGGTGCC

GTAGCGCTGAAGTCTTACGAGGAAGAGTTGGCGAAAGATCCACGTATTGCCGCCACCATGGAAAACGCC

CAGAAAGGTGAAATCATGCCGAACATCCCGCAGATGTCCGCTTTCTGGTATGCCGTGCGTACTGCGGTG

ATCAACGCCGCCAGCGGTCGTCAGACTGTCGATGAAGCCCTGAAAGACGCGCAGACTCGTATCACCAAG

TAA

>MBP_cpEGFP208 amino acid

MKIEEGKLVIWINGDKGYNGLAEVGKKFEKDTGIKVTVEHPDKLEEKFPQVAATGDGPDIIFWAHDRFG

GYAQSGLLAEITPDKAFQDKLYPFTWDAVRYNGKLIAYPIAVEALSLIYNKDLLPNPPKTWEEIPALDK

ELKAKGKSALMFNLQEPYFTWPLIAADGGYAFKYENGKYDIKDVGVDNAGAKAGLTFLVDLIKNKHMNA

DTDASYNVFIMADKQKNGIKANFKIRHNIEDGGVQLAYHYQQNTPIGDGPVLLPDNHYLSVQSKLSKDP

NEKRDHMVLLEFVTAAGITLGMDELYKGGTGGSMVSKGEELFTGVVPILVELDGDVNGHKFSVSGEGEG

DATYGKLTLKFICTTGKLPVPWPTLVTTLTYGVQCFSRYPDHMKQHDFFKSAMPEGYIQERTIFFKDDG

NYKTRAEVKFEGDTLVNRIELKGIDFKEDGNILGHKLEYNFNASTDYSIAEAAFNKGETAMTINGPWAW

SNIDTSKVNYGVTVLPTFKGQPSKPFVGVLSAGINAASPNKELAKEFLENYLLTDEGLEAVNKDKPLGA

VALKSYEEELAKDPRIAATMENAQKGEIMPNIPQMSAFWYAVRTAVINAASGRQTVDEALKDAQTRITK

*

>MBP_cpEGFP211 nucleotide

ATGAAAATCGAAGAAGGTAAACTGGTAATCTGGATTAACGGCGATAAAGGCTATAACGGACTCGCTGAA

GTCGGTAAGAAATTCGAGAAAGATACCGGAATTAAAGTCACCGTTGAGCATCCGGATAAACTGGAAGAG

AAATTCCCACAGGTTGCGGCAACTGGCGATGGCCCTGACATTATCTTCTGGGCACACGACCGCTTTGGT

GGCTACGCTCAATCTGGCCTGTTGGCTGAAATCACCCCGGACAAAGCGTTCCAGGACAAGCTGTATCCG

TTTACCTGGGATGCCGTACGTTACAACGGCAAGCTGATTGCTTACCCGATCGCTGTTGAAGCGTTATCG

CTGATTTATAACAAAGATCTGCTGCCGAACCCGCCAAAAACCTGGGAAGAGATCCCGGCGCTGGATAAA

GAACTGAAAGCGAAAGGTAAGAGCGCGCTGATGTTCAACCTGCAAGAACCGTACTTCACCTGGCCGCTG

ATTGCTGCTGACGGGGGTTATGCGTTCAAGTATGAAAACGGCAAGTACGACATTAAAGACGTGGGCGTG

GATAACGCTGGCGCGAAAGCGGGTCTGACCTTCCTGGTTGACCTGATTAAAAACAAACACATGAATGCA

GACACCGATTACTCCATCGCATCTTATAACGTCTTTATCATGGCCGACAAGCAGAAGAACGGCATCAAG

GCGAACTTCAAGATCCGCCACAACATCGAGGACGGCGGCGTGCAGCTCGCCTATCACTACCAGCAGAAC

ACCCCCATCGGCGACGGCCCCGTGCTGCTGCCCGACAACCACTACCTGAGCGTGCAGTCCAAACTGAGC

AAAGACCCCAACGAGAAGCGCGATCACATGGTCCTGCTGGAGTTCGTGACCGCCGCCGGGATCACTCTC

GGCATGGACGAGCTGTACAAGGGCGGTACCGGAGGGAGCATGGTGAGCAAGGGCGAGGAGCTGTTCACC

GGGGTGGTGCCCATCCTGGTCGAGCTGGACGGCGACGTAAACGGCCACAAGTTCAGCGTGTCCGGCGAG

GGCGAGGGCGATGCCACCTACGGCAAGCTGACCCTGAAGTTCATCTGCACCACCGGCAAGCTGCCCGTG

CCCTGGCCCACCCTCGTGACCACCCTGACCTACGGCGTGCAGTGCTTCAGCCGCTACCCCGACCACATG

AAGCAGCACGACTTCTTCAAGTCCGCCATGCCCGAAGGCTACATTCAGGAGCGCACCATCTTCTTCAAG

GACGACGGCAACTATAAGACACGCGCTGAGGTTAAGTTCGAGGGCGACACTCTGGTTAACCGCATCGAG

CTGAAGGGCATCGACTTCAAGGAGGACGGCAACATCCTGGGCCATAAGCTTGAATATAACTTCAACGCA

TCTTCCATCGCAGAAGCTGCCTTTAATAAAGGCGAAACAGCGATGACCATCAACGGCCCGTGGGCATGG

TCCAACATCGACACCAGCAAAGTGAATTATGGTGTAACGGTACTGCCGACCTTCAAGGGTCAACCATCC

AAACCGTTCGTTGGCGTGCTGAGCGCAGGTATTAACGCCGCCAGTCCGAACAAAGAGCTGGCGAAAGAG

TTCCTCGAAAACTATCTGCTGACTGATGAAGGTCTGGAAGCGGTTAATAAAGACAAACCGCTGGGTGCC

GTAGCGCTGAAGTCTTACGAGGAAGAGTTGGCGAAAGATCCACGTATTGCCGCCACCATGGAAAACGCC

CAGAAAGGTGAAATCATGCCGAACATCCCGCAGATGTCCGCTTTCTGGTATGCCGTGCGTACTGCGGTG

ATCAACGCCGCCAGCGGTCGTCAGACTGTCGATGAAGCCCTGAAAGACGCGCAGACTCGTATCACCAAG

TAA

>MBP_cpEGFP208 amino acid

MKIEEGKLVIWINGDKGYNGLAEVGKKFEKDTGIKVTVEHPDKLEEKFPQVAATGDGPDIIFWAHDRFG

GYAQSGLLAEITPDKAFQDKLYPFTWDAVRYNGKLIAYPIAVEALSLIYNKDLLPNPPKTWEEIPALDK

ELKAKGKSALMFNLQEPYFTWPLIAADGGYAFKYENGKYDIKDVGVDNAGAKAGLTFLVDLIKNKHMNA

DTDYSIASYNVFIMADKQKNGIKANFKIRHNIEDGGVQLAYHYQQNTPIGDGPVLLPDNHYLSVQSKLS

KDPNEKRDHMVLLEFVTAAGITLGMDELYKGGTGGSMVSKGEELFTGVVPILVELDGDVNGHKFSVSGE

GEGDATYGKLTLKFICTTGKLPVPWPTLVTTLTYGVQCFSRYPDHMKQHDFFKSAMPEGYIQERTIFFK

DDGNYKTRAEVKFEGDTLVNRIELKGIDFKEDGNILGHKLEYNFNASSIAEAAFNKGETAMTINGPWAW

SNIDTSKVNYGVTVLPTFKGQPSKPFVGVLSAGINAASPNKELAKEFLENYLLTDEGLEAVNKDKPLGA

VALKSYEEELAKDPRIAATMENAQKGEIMPNIPQMSAFWYAVRTAVINAASGRQTVDEALKDAQTRITK

*

>MBP_cpEGFP214 nucleotide

ATGAAAATCGAAGAAGGTAAACTGGTAATCTGGATTAACGGCGATAAAGGCTATAACGGACTCGCTGAA

GTCGGTAAGAAATTCGAGAAAGATACCGGAATTAAAGTCACCGTTGAGCATCCGGATAAACTGGAAGAG

AAATTCCCACAGGTTGCGGCAACTGGCGATGGCCCTGACATTATCTTCTGGGCACACGACCGCTTTGGT

GGCTACGCTCAATCTGGCCTGTTGGCTGAAATCACCCCGGACAAAGCGTTCCAGGACAAGCTGTATCCG

TTTACCTGGGATGCCGTACGTTACAACGGCAAGCTGATTGCTTACCCGATCGCTGTTGAAGCGTTATCG

CTGATTTATAACAAAGATCTGCTGCCGAACCCGCCAAAAACCTGGGAAGAGATCCCGGCGCTGGATAAA

GAACTGAAAGCGAAAGGTAAGAGCGCGCTGATGTTCAACCTGCAAGAACCGTACTTCACCTGGCCGCTG

ATTGCTGCTGACGGGGGTTATGCGTTCAAGTATGAAAACGGCAAGTACGACATTAAAGACGTGGGCGTG

GATAACGCTGGCGCGAAAGCGGGTCTGACCTTCCTGGTTGACCTGATTAAAAACAAACACATGAATGCA

GACACCGATTACTCCATCGCAGAAGCTGCATCTTATAACGTCTTTATCATGGCCGACAAGCAGAAGAAC

GGCATCAAGGCGAACTTCAAGATCCGCCACAACATCGAGGACGGCGGCGTGCAGCTCGCCTATCACTAC

CAGCAGAACACCCCCATCGGCGACGGCCCCGTGCTGCTGCCCGACAACCACTACCTGAGCGTGCAGTCC

AAACTGAGCAAAGACCCCAACGAGAAGCGCGATCACATGGTCCTGCTGGAGTTCGTGACCGCCGCCGGG

ATCACTCTCGGCATGGACGAGCTGTACAAGGGCGGTACCGGAGGGAGCATGGTGAGCAAGGGCGAGGAG

CTGTTCACCGGGGTGGTGCCCATCCTGGTCGAGCTGGACGGCGACGTAAACGGCCACAAGTTCAGCGTG

TCCGGCGAGGGCGAGGGCGATGCCACCTACGGCAAGCTGACCCTGAAGTTCATCTGCACCACCGGCAAG

CTGCCCGTGCCCTGGCCCACCCTCGTGACCACCCTGACCTACGGCGTGCAGTGCTTCAGCCGCTACCCC

GACCACATGAAGCAGCACGACTTCTTCAAGTCCGCCATGCCCGAAGGCTACATTCAGGAGCGCACCATC

TTCTTCAAGGACGACGGCAACTATAAGACACGCGCTGAGGTTAAGTTCGAGGGCGACACTCTGGTTAAC

CGCATCGAGCTGAAGGGCATCGACTTCAAGGAGGACGGCAACATCCTGGGCCATAAGCTTGAATATAAC

TTCAACGCATCTGAAGCTGCCTTTAATAAAGGCGAAACAGCGATGACCATCAACGGCCCGTGGGCATGG

TCCAACATCGACACCAGCAAAGTGAATTATGGTGTAACGGTACTGCCGACCTTCAAGGGTCAACCATCC

AAACCGTTCGTTGGCGTGCTGAGCGCAGGTATTAACGCCGCCAGTCCGAACAAAGAGCTGGCGAAAGAG

TTCCTCGAAAACTATCTGCTGACTGATGAAGGTCTGGAAGCGGTTAATAAAGACAAACCGCTGGGTGCC

GTAGCGCTGAAGTCTTACGAGGAAGAGTTGGCGAAAGATCCACGTATTGCCGCCACCATGGAAAACGCC

CAGAAAGGTGAAATCATGCCGAACATCCCGCAGATGTCCGCTTTCTGGTATGCCGTGCGTACTGCGGTG

ATCAACGCCGCCAGCGGTCGTCAGACTGTCGATGAAGCCCTGAAAGACGCGCAGACTCGTATCACCAAG

TAA

>MBP_cpEGFP214 amino acid

MKIEEGKLVIWINGDKGYNGLAEVGKKFEKDTGIKVTVEHPDKLEEKFPQVAATGDGPDIIFWAHDRFG

GYAQSGLLAEITPDKAFQDKLYPFTWDAVRYNGKLIAYPIAVEALSLIYNKDLLPNPPKTWEEIPALDK

ELKAKGKSALMFNLQEPYFTWPLIAADGGYAFKYENGKYDIKDVGVDNAGAKAGLTFLVDLIKNKHMNA

DTDYSIAEAASYNVFIMADKQKNGIKANFKIRHNIEDGGVQLAYHYQQNTPIGDGPVLLPDNHYLSVQS

KLSKDPNEKRDHMVLLEFVTAAGITLGMDELYKGGTGGSMVSKGEELFTGVVPILVELDGDVNGHKFSV

SGEGEGDATYGKLTLKFICTTGKLPVPWPTLVTTLTYGVQCFSRYPDHMKQHDFFKSAMPEGYIQERTI

FFKDDGNYKTRAEVKFEGDTLVNRIELKGIDFKEDGNILGHKLEYNFNASEAAFNKGETAMTINGPWAW

SNIDTSKVNYGVTVLPTFKGQPSKPFVGVLSAGINAASPNKELAKEFLENYLLTDEGLEAVNKDKPLGA

VALKSYEEELAKDPRIAATMENAQKGEIMPNIPQMSAFWYAVRTAVINAASGRQTVDEALKDAQTRITK

*

>MBP_cpEGFP217 nucleotide

ATGAAAATCGAAGAAGGTAAACTGGTAATCTGGATTAACGGCGATAAAGGCTATAACGGACTCGCTGAA

GTCGGTAAGAAATTCGAGAAAGATACCGGAATTAAAGTCACCGTTGAGCATCCGGATAAACTGGAAGAG

AAATTCCCACAGGTTGCGGCAACTGGCGATGGCCCTGACATTATCTTCTGGGCACACGACCGCTTTGGT

GGCTACGCTCAATCTGGCCTGTTGGCTGAAATCACCCCGGACAAAGCGTTCCAGGACAAGCTGTATCCG

TTTACCTGGGATGCCGTACGTTACAACGGCAAGCTGATTGCTTACCCGATCGCTGTTGAAGCGTTATCG

CTGATTTATAACAAAGATCTGCTGCCGAACCCGCCAAAAACCTGGGAAGAGATCCCGGCGCTGGATAAA

GAACTGAAAGCGAAAGGTAAGAGCGCGCTGATGTTCAACCTGCAAGAACCGTACTTCACCTGGCCGCTG

ATTGCTGCTGACGGGGGTTATGCGTTCAAGTATGAAAACGGCAAGTACGACATTAAAGACGTGGGCGTG

GATAACGCTGGCGCGAAAGCGGGTCTGACCTTCCTGGTTGACCTGATTAAAAACAAACACATGAATGCA

GACACCGATTACTCCATCGCAGAAGCTGCCTTTAATGCATCTTATAACGTCTTTATCATGGCCGACAAG

CAGAAGAACGGCATCAAGGCGAACTTCAAGATCCGCCACAACATCGAGGACGGCGGCGTGCAGCTCGCC

TATCACTACCAGCAGAACACCCCCATCGGCGACGGCCCCGTGCTGCTGCCCGACAACCACTACCTGAGC

GTGCAGTCCAAACTGAGCAAAGACCCCAACGAGAAGCGCGATCACATGGTCCTGCTGGAGTTCGTGACC

GCCGCCGGGATCACTCTCGGCATGGACGAGCTGTACAAGGGCGGTACCGGAGGGAGCATGGTGAGCAAG

GGCGAGGAGCTGTTCACCGGGGTGGTGCCCATCCTGGTCGAGCTGGACGGCGACGTAAACGGCCACAAG

TTCAGCGTGTCCGGCGAGGGCGAGGGCGATGCCACCTACGGCAAGCTGACCCTGAAGTTCATCTGCACC

ACCGGCAAGCTGCCCGTGCCCTGGCCCACCCTCGTGACCACCCTGACCTACGGCGTGCAGTGCTTCAGC

CGCTACCCCGACCACATGAAGCAGCACGACTTCTTCAAGTCCGCCATGCCCGAAGGCTACATTCAGGAG

CGCACCATCTTCTTCAAGGACGACGGCAACTATAAGACACGCGCTGAGGTTAAGTTCGAGGGCGACACT

CTGGTTAACCGCATCGAGCTGAAGGGCATCGACTTCAAGGAGGACGGCAACATCCTGGGCCATAAGCTT

GAATATAACTTCAACGCATCTTTTAATAAAGGCGAAACAGCGATGACCATCAACGGCCCGTGGGCATGG

TCCAACATCGACACCAGCAAAGTGAATTATGGTGTAACGGTACTGCCGACCTTCAAGGGTCAACCATCC

AAACCGTTCGTTGGCGTGCTGAGCGCAGGTATTAACGCCGCCAGTCCGAACAAAGAGCTGGCGAAAGAG

TTCCTCGAAAACTATCTGCTGACTGATGAAGGTCTGGAAGCGGTTAATAAAGACAAACCGCTGGGTGCC

GTAGCGCTGAAGTCTTACGAGGAAGAGTTGGCGAAAGATCCACGTATTGCCGCCACCATGGAAAACGCC

CAGAAAGGTGAAATCATGCCGAACATCCCGCAGATGTCCGCTTTCTGGTATGCCGTGCGTACTGCGGTG

ATCAACGCCGCCAGCGGTCGTCAGACTGTCGATGAAGCCCTGAAAGACGCGCAGACTCGTATCACCAAG

TAA

>MBP_cpEGFP217 amino acid MKIEEGKLVIWINGDKGYNGLAEVGKKFEKDTGIKVTVEHPDKLEEKFPQVAATGDGPDIIFWAHDRFG

GYAQSGLLAEITPDKAFQDKLYPFTWDAVRYNGKLIAYPIAVEALSLIYNKDLLPNPPKTWEEIPALDK

ELKAKGKSALMFNLQEPYFTWPLIAADGGYAFKYENGKYDIKDVGVDNAGAKAGLTFLVDLIKNKHMNA

DTDYSIAEAAFNASYNVFIMADKQKNGIKANFKIRHNIEDGGVQLAYHYQQNTPIGDGPVLLPDNHYLS

VQSKLSKDPNEKRDHMVLLEFVTAAGITLGMDELYKGGTGGSMVSKGEELFTGVVPILVELDGDVNGHK

FSVSGEGEGDATYGKLTLKFICTTGKLPVPWPTLVTTLTYGVQCFSRYPDHMKQHDFFKSAMPEGYIQE

RTIFFKDDGNYKTRAEVKFEGDTLVNRIELKGIDFKEDGNILGHKLEYNFNASFNKGETAMTINGPWAW

SNIDTSKVNYGVTVLPTFKGQPSKPFVGVLSAGINAASPNKELAKEFLENYLLTDEGLEAVNKDKPLGA

VALKSYEEELAKDPRIAATMENAQKGEIMPNIPQMSAFWYAVRTAVINAASGRQTVDEALKDAQTRITK

*

>MBP_cpEGFP224 nucleotide

ATGAAAATCGAAGAAGGTAAACTGGTAATCTGGATTAACGGCGATAAAGGCTATAACGGACTCGCTGAA

GTCGGTAAGAAATTCGAGAAAGATACCGGAATTAAAGTCACCGTTGAGCATCCGGATAAACTGGAAGAG

AAATTCCCACAGGTTGCGGCAACTGGCGATGGCCCTGACATTATCTTCTGGGCACACGACCGCTTTGGT

GGCTACGCTCAATCTGGCCTGTTGGCTGAAATCACCCCGGACAAAGCGTTCCAGGACAAGCTGTATCCG

TTTACCTGGGATGCCGTACGTTACAACGGCAAGCTGATTGCTTACCCGATCGCTGTTGAAGCGTTATCG

CTGATTTATAACAAAGATCTGCTGCCGAACCCGCCAAAAACCTGGGAAGAGATCCCGGCGCTGGATAAA

GAACTGAAAGCGAAAGGTAAGAGCGCGCTGATGTTCAACCTGCAAGAACCGTACTTCACCTGGCCGCTG

ATTGCTGCTGACGGGGGTTATGCGTTCAAGTATGAAAACGGCAAGTACGACATTAAAGACGTGGGCGTG

GATAACGCTGGCGCGAAAGCGGGTCTGACCTTCCTGGTTGACCTGATTAAAAACAAACACATGAATGCA

GACACCGATTACTCCATCGCAGAAGCTGCCTTTAATAAAGGCGAAACAGCGATGACCGCATCTTATAAC

GTCTTTATCATGGCCGACAAGCAGAAGAACGGCATCAAGGCGAACTTCAAGATCCGCCACAACATCGAG

GACGGCGGCGTGCAGCTCGCCTATCACTACCAGCAGAACACCCCCATCGGCGACGGCCCCGTGCTGCTG

CCCGACAACCACTACCTGAGCGTGCAGTCCAAACTGAGCAAAGACCCCAACGAGAAGCGCGATCACATG

GTCCTGCTGGAGTTCGTGACCGCCGCCGGGATCACTCTCGGCATGGACGAGCTGTACAAGGGCGGTACC

GGAGGGAGCATGGTGAGCAAGGGCGAGGAGCTGTTCACCGGGGTGGTGCCCATCCTGGTCGAGCTGGAC

GGCGACGTAAACGGCCACAAGTTCAGCGTGTCCGGCGAGGGCGAGGGCGATGCCACCTACGGCAAGCTG

ACCCTGAAGTTCATCTGCACCACCGGCAAGCTGCCCGTGCCCTGGCCCACCCTCGTGACCACCCTGACC

TACGGCGTGCAGTGCTTCAGCCGCTACCCCGACCACATGAAGCAGCACGACTTCTTCAAGTCCGCCATG

CCCGAAGGCTACATTCAGGAGCGCACCATCTTCTTCAAGGACGACGGCAACTATAAGACACGCGCTGAG

GTTAAGTTCGAGGGCGACACTCTGGTTAACCGCATCGAGCTGAAGGGCATCGACTTCAAGGAGGACGGC

AACATCCTGGGCCATAAGCTTGAATATAACTTCAACGCATCTATGACCATCAACGGCCCGTGGGCATGG

TCCAACATCGACACCAGCAAAGTGAATTATGGTGTAACGGTACTGCCGACCTTCAAGGGTCAACCATCC

AAACCGTTCGTTGGCGTGCTGAGCGCAGGTATTAACGCCGCCAGTCCGAACAAAGAGCTGGCGAAAGAG

TTCCTCGAAAACTATCTGCTGACTGATGAAGGTCTGGAAGCGGTTAATAAAGACAAACCGCTGGGTGCC

GTAGCGCTGAAGTCTTACGAGGAAGAGTTGGCGAAAGATCCACGTATTGCCGCCACCATGGAAAACGCC

CAGAAAGGTGAAATCATGCCGAACATCCCGCAGATGTCCGCTTTCTGGTATGCCGTGCGTACTGCGGTG

ATCAACGCCGCCAGCGGTCGTCAGACTGTCGATGAAGCCCTGAAAGACGCGCAGACTCGTATCACCAAG

TAA

>MBP_cpEGFP224 amino acid

MKIEEGKLVIWINGDKGYNGLAEVGKKFEKDTGIKVTVEHPDKLEEKFPQVAATGDGPDIIFWAHDRFG

GYAQSGLLAEITPDKAFQDKLYPFTWDAVRYNGKLIAYPIAVEALSLIYNKDLLPNPPKTWEEIPALDK

ELKAKGKSALMFNLQEPYFTWPLIAADGGYAFKYENGKYDIKDVGVDNAGAKAGLTFLVDLIKNKHMNA

DTDYSIAEAAFNKGETAMTASYNVFIMADKQKNGIKANFKIRHNIEDGGVQLAYHYQQNTPIGDGPVLL

PDNHYLSVQSKLSKDPNEKRDHMVLLEFVTAAGITLGMDELYKGGTGGSMVSKGEELFTGVVPILVELD

GDVNGHKFSVSGEGEGDATYGKLTLKFICTTGKLPVPWPTLVTTLTYGVQCFSRYPDHMKQHDFFKSAM

PEGYIQERTIFFKDDGNYKTRAEVKFEGDTLVNRIELKGIDFKEDGNILGHKLEYNFNASMTINGPWAW

SNIDTSKVNYGVTVLPTFKGQPSKPFVGVLSAGINAASPNKELAKEFLENYLLTDEGLEAVNKDKPLGA

VALKSYEEELAKDPRIAATMENAQKGEIMPNIPQMSAFWYAVRTAVINAASGRQTVDEALKDAQTRITK

*

>MBP_cpEGFP227 nucleotide

ATGAAAATCGAAGAAGGTAAACTGGTAATCTGGATTAACGGCGATAAAGGCTATAACGGACTCGCTGAA

GTCGGTAAGAAATTCGAGAAAGATACCGGAATTAAAGTCACCGTTGAGCATCCGGATAAACTGGAAGAG

AAATTCCCACAGGTTGCGGCAACTGGCGATGGCCCTGACATTATCTTCTGGGCACACGACCGCTTTGGT

GGCTACGCTCAATCTGGCCTGTTGGCTGAAATCACCCCGGACAAAGCGTTCCAGGACAAGCTGTATCCG

TTTACCTGGGATGCCGTACGTTACAACGGCAAGCTGATTGCTTACCCGATCGCTGTTGAAGCGTTATCG

CTGATTTATAACAAAGATCTGCTGCCGAACCCGCCAAAAACCTGGGAAGAGATCCCGGCGCTGGATAAA

GAACTGAAAGCGAAAGGTAAGAGCGCGCTGATGTTCAACCTGCAAGAACCGTACTTCACCTGGCCGCTG

ATTGCTGCTGACGGGGGTTATGCGTTCAAGTATGAAAACGGCAAGTACGACATTAAAGACGTGGGCGTG

GATAACGCTGGCGCGAAAGCGGGTCTGACCTTCCTGGTTGACCTGATTAAAAACAAACACATGAATGCA

GACACCGATTACTCCATCGCAGAAGCTGCCTTTAATAAAGGCGAAACAGCGATGACCATCAACGGCGCA

TCTTATAACGTCTTTATCATGGCCGACAAGCAGAAGAACGGCATCAAGGCGAACTTCAAGATCCGCCAC

AACATCGAGGACGGCGGCGTGCAGCTCGCCTATCACTACCAGCAGAACACCCCCATCGGCGACGGCCCC

GTGCTGCTGCCCGACAACCACTACCTGAGCGTGCAGTCCAAACTGAGCAAAGACCCCAACGAGAAGCGC

GATCACATGGTCCTGCTGGAGTTCGTGACCGCCGCCGGGATCACTCTCGGCATGGACGAGCTGTACAAG

GGCGGTACCGGAGGGAGCATGGTGAGCAAGGGCGAGGAGCTGTTCACCGGGGTGGTGCCCATCCTGGTC

GAGCTGGACGGCGACGTAAACGGCCACAAGTTCAGCGTGTCCGGCGAGGGCGAGGGCGATGCCACCTAC

GGCAAGCTGACCCTGAAGTTCATCTGCACCACCGGCAAGCTGCCCGTGCCCTGGCCCACCCTCGTGACC

ACCCTGACCTACGGCGTGCAGTGCTTCAGCCGCTACCCCGACCACATGAAGCAGCACGACTTCTTCAAG

TCCGCCATGCCCGAAGGCTACATTCAGGAGCGCACCATCTTCTTCAAGGACGACGGCAACTATAAGACA

CGCGCTGAGGTTAAGTTCGAGGGCGACACTCTGGTTAACCGCATCGAGCTGAAGGGCATCGACTTCAAG

GAGGACGGCAACATCCTGGGCCATAAGCTTGAATATAACTTCAACGCATCTAACGGCCCGTGGGCATGG

TCCAACATCGACACCAGCAAAGTGAATTATGGTGTAACGGTACTGCCGACCTTCAAGGGTCAACCATCC

AAACCGTTCGTTGGCGTGCTGAGCGCAGGTATTAACGCCGCCAGTCCGAACAAAGAGCTGGCGAAAGAG

TTCCTCGAAAACTATCTGCTGACTGATGAAGGTCTGGAAGCGGTTAATAAAGACAAACCGCTGGGTGCC

GTAGCGCTGAAGTCTTACGAGGAAGAGTTGGCGAAAGATCCACGTATTGCCGCCACCATGGAAAACGCC

CAGAAAGGTGAAATCATGCCGAACATCCCGCAGATGTCCGCTTTCTGGTATGCCGTGCGTACTGCGGTG

ATCAACGCCGCCAGCGGTCGTCAGACTGTCGATGAAGCCCTGAAAGACGCGCAGACTCGTATCACCAAG

TAA

>MBP_cpEGFP227 amino acid

MKIEEGKLVIWINGDKGYNGLAEVGKKFEKDTGIKVTVEHPDKLEEKFPQVAATGDGPDIIFWAHDRFG

GYAQSGLLAEITPDKAFQDKLYPFTWDAVRYNGKLIAYPIAVEALSLIYNKDLLPNPPKTWEEIPALDK

ELKAKGKSALMFNLQEPYFTWPLIAADGGYAFKYENGKYDIKDVGVDNAGAKAGLTFLVDLIKNKHMNA

DTDYSIAEAAFNKGETAMTINGASYNVFIMADKQKNGIKANFKIRHNIEDGGVQLAYHYQQNTPIGDGP

VLLPDNHYLSVQSKLSKDPNEKRDHMVLLEFVTAAGITLGMDELYKGGTGGSMVSKGEELFTGVVPILV

ELDGDVNGHKFSVSGEGEGDATYGKLTLKFICTTGKLPVPWPTLVTTLTYGVQCFSRYPDHMKQHDFFK

SAMPEGYIQERTIFFKDDGNYKTRAEVKFEGDTLVNRIELKGIDFKEDGNILGHKLEYNFNASNGPWAW

SNIDTSKVNYGVTVLPTFKGQPSKPFVGVLSAGINAASPNKELAKEFLENYLLTDEGLEAVNKDKPLGA

VALKSYEEELAKDPRIAATMENAQKGEIMPNIPQMSAFWYAVRTAVINAASGRQTVDEALKDAQTRITK

*

>MBP_cpEGFP230 nucleotide

ATGAAAATCGAAGAAGGTAAACTGGTAATCTGGATTAACGGCGATAAAGGCTATAACGGACTCGCTGAA

GTCGGTAAGAAATTCGAGAAAGATACCGGAATTAAAGTCACCGTTGAGCATCCGGATAAACTGGAAGAG

AAATTCCCACAGGTTGCGGCAACTGGCGATGGCCCTGACATTATCTTCTGGGCACACGACCGCTTTGGT

GGCTACGCTCAATCTGGCCTGTTGGCTGAAATCACCCCGGACAAAGCGTTCCAGGACAAGCTGTATCCG

TTTACCTGGGATGCCGTACGTTACAACGGCAAGCTGATTGCTTACCCGATCGCTGTTGAAGCGTTATCG

CTGATTTATAACAAAGATCTGCTGCCGAACCCGCCAAAAACCTGGGAAGAGATCCCGGCGCTGGATAAA

GAACTGAAAGCGAAAGGTAAGAGCGCGCTGATGTTCAACCTGCAAGAACCGTACTTCACCTGGCCGCTG

ATTGCTGCTGACGGGGGTTATGCGTTCAAGTATGAAAACGGCAAGTACGACATTAAAGACGTGGGCGTG

GATAACGCTGGCGCGAAAGCGGGTCTGACCTTCCTGGTTGACCTGATTAAAAACAAACACATGAATGCA

GACACCGATTACTCCATCGCAGAAGCTGCCTTTAATAAAGGCGAAACAGCGATGACCATCAACGGCCCG

TGGGCAGCATCTTATAACGTCTTTATCATGGCCGACAAGCAGAAGAACGGCATCAAGGCGAACTTCAAG

ATCCGCCACAACATCGAGGACGGCGGCGTGCAGCTCGCCTATCACTACCAGCAGAACACCCCCATCGGC

GACGGCCCCGTGCTGCTGCCCGACAACCACTACCTGAGCGTGCAGTCCAAACTGAGCAAAGACCCCAAC

GAGAAGCGCGATCACATGGTCCTGCTGGAGTTCGTGACCGCCGCCGGGATCACTCTCGGCATGGACGAG

CTGTACAAGGGCGGTACCGGAGGGAGCATGGTGAGCAAGGGCGAGGAGCTGTTCACCGGGGTGGTGCCC

ATCCTGGTCGAGCTGGACGGCGACGTAAACGGCCACAAGTTCAGCGTGTCCGGCGAGGGCGAGGGCGAT

GCCACCTACGGCAAGCTGACCCTGAAGTTCATCTGCACCACCGGCAAGCTGCCCGTGCCCTGGCCCACC

CTCGTGACCACCCTGACCTACGGCGTGCAGTGCTTCAGCCGCTACCCCGACCACATGAAGCAGCACGAC

TTCTTCAAGTCCGCCATGCCCGAAGGCTACATTCAGGAGCGCACCATCTTCTTCAAGGACGACGGCAAC

TATAAGACACGCGCTGAGGTTAAGTTCGAGGGCGACACTCTGGTTAACCGCATCGAGCTGAAGGGCATC

GACTTCAAGGAGGACGGCAACATCCTGGGCCATAAGCTTGAATATAACTTCAACGCATCTTGGGCATGG

TCCAACATCGACACCAGCAAAGTGAATTATGGTGTAACGGTACTGCCGACCTTCAAGGGTCAACCATCC

AAACCGTTCGTTGGCGTGCTGAGCGCAGGTATTAACGCCGCCAGTCCGAACAAAGAGCTGGCGAAAGAG

TTCCTCGAAAACTATCTGCTGACTGATGAAGGTCTGGAAGCGGTTAATAAAGACAAACCGCTGGGTGCC

GTAGCGCTGAAGTCTTACGAGGAAGAGTTGGCGAAAGATCCACGTATTGCCGCCACCATGGAAAACGCC

CAGAAAGGTGAAATCATGCCGAACATCCCGCAGATGTCCGCTTTCTGGTATGCCGTGCGTACTGCGGTG

ATCAACGCCGCCAGCGGTCGTCAGACTGTCGATGAAGCCCTGAAAGACGCGCAGACTCGTATCACCAAG

TAA

>MBP_cpEGFP230 amino acid

MKIEEGKLVIWINGDKGYNGLAEVGKKFEKDTGIKVTVEHPDKLEEKFPQVAATGDGPDIIFWAHDRFG

GYAQSGLLAEITPDKAFQDKLYPFTWDAVRYNGKLIAYPIAVEALSLIYNKDLLPNPPKTWEEIPALDK

ELKAKGKSALMFNLQEPYFTWPLIAADGGYAFKYENGKYDIKDVGVDNAGAKAGLTFLVDLIKNKHMNA

DTDYSIAEAAFNKGETAMTINGPWAASYNVFIMADKQKNGIKANFKIRHNIEDGGVQLAYHYQQNTPIG

DGPVLLPDNHYLSVQSKLSKDPNEKRDHMVLLEFVTAAGITLGMDELYKGGTGGSMVSKGEELFTGVVP

ILVELDGDVNGHKFSVSGEGEGDATYGKLTLKFICTTGKLPVPWPTLVTTLTYGVQCFSRYPDHMKQHD

FFKSAMPEGYIQERTIFFKDDGNYKTRAEVKFEGDTLVNRIELKGIDFKEDGNILGHKLEYNFNASWAW

SNIDTSKVNYGVTVLPTFKGQPSKPFVGVLSAGINAASPNKELAKEFLENYLLTDEGLEAVNKDKPLGA

VALKSYEEELAKDPRIAATMENAQKGEIMPNIPQMSAFWYAVRTAVINAASGRQTVDEALKDAQTRITK

*

>MBP_cpEGFP233 nucleotide

ATGAAAATCGAAGAAGGTAAACTGGTAATCTGGATTAACGGCGATAAAGGCTATAACGGACTCGCTGAA

GTCGGTAAGAAATTCGAGAAAGATACCGGAATTAAAGTCACCGTTGAGCATCCGGATAAACTGGAAGAG

AAATTCCCACAGGTTGCGGCAACTGGCGATGGCCCTGACATTATCTTCTGGGCACACGACCGCTTTGGT

GGCTACGCTCAATCTGGCCTGTTGGCTGAAATCACCCCGGACAAAGCGTTCCAGGACAAGCTGTATCCG

TTTACCTGGGATGCCGTACGTTACAACGGCAAGCTGATTGCTTACCCGATCGCTGTTGAAGCGTTATCG

CTGATTTATAACAAAGATCTGCTGCCGAACCCGCCAAAAACCTGGGAAGAGATCCCGGCGCTGGATAAA

GAACTGAAAGCGAAAGGTAAGAGCGCGCTGATGTTCAACCTGCAAGAACCGTACTTCACCTGGCCGCTG

ATTGCTGCTGACGGGGGTTATGCGTTCAAGTATGAAAACGGCAAGTACGACATTAAAGACGTGGGCGTG

GATAACGCTGGCGCGAAAGCGGGTCTGACCTTCCTGGTTGACCTGATTAAAAACAAACACATGAATGCA

GACACCGATTACTCCATCGCAGAAGCTGCCTTTAATAAAGGCGAAACAGCGATGACCATCAACGGCCCG

TGGGCATGGTCCAACGCATCTTATAACGTCTTTATCATGGCCGACAAGCAGAAGAACGGCATCAAGGCG

AACTTCAAGATCCGCCACAACATCGAGGACGGCGGCGTGCAGCTCGCCTATCACTACCAGCAGAACACC

CCCATCGGCGACGGCCCCGTGCTGCTGCCCGACAACCACTACCTGAGCGTGCAGTCCAAACTGAGCAAA

GACCCCAACGAGAAGCGCGATCACATGGTCCTGCTGGAGTTCGTGACCGCCGCCGGGATCACTCTCGGC

ATGGACGAGCTGTACAAGGGCGGTACCGGAGGGAGCATGGTGAGCAAGGGCGAGGAGCTGTTCACCGGG

GTGGTGCCCATCCTGGTCGAGCTGGACGGCGACGTAAACGGCCACAAGTTCAGCGTGTCCGGCGAGGGC

GAGGGCGATGCCACCTACGGCAAGCTGACCCTGAAGTTCATCTGCACCACCGGCAAGCTGCCCGTGCCC

TGGCCCACCCTCGTGACCACCCTGACCTACGGCGTGCAGTGCTTCAGCCGCTACCCCGACCACATGAAG

CAGCACGACTTCTTCAAGTCCGCCATGCCCGAAGGCTACATTCAGGAGCGCACCATCTTCTTCAAGGAC

GACGGCAACTATAAGACACGCGCTGAGGTTAAGTTCGAGGGCGACACTCTGGTTAACCGCATCGAGCTG

AAGGGCATCGACTTCAAGGAGGACGGCAACATCCTGGGCCATAAGCTTGAATATAACTTCAACGCATCT

TCCAACATCGACACCAGCAAAGTGAATTATGGTGTAACGGTACTGCCGACCTTCAAGGGTCAACCATCC

AAACCGTTCGTTGGCGTGCTGAGCGCAGGTATTAACGCCGCCAGTCCGAACAAAGAGCTGGCGAAAGAG

TTCCTCGAAAACTATCTGCTGACTGATGAAGGTCTGGAAGCGGTTAATAAAGACAAACCGCTGGGTGCC

GTAGCGCTGAAGTCTTACGAGGAAGAGTTGGCGAAAGATCCACGTATTGCCGCCACCATGGAAAACGCC

CAGAAAGGTGAAATCATGCCGAACATCCCGCAGATGTCCGCTTTCTGGTATGCCGTGCGTACTGCGGTG

ATCAACGCCGCCAGCGGTCGTCAGACTGTCGATGAAGCCCTGAAAGACGCGCAGACTCGTATCACCAAG

TAA

>MBP_cpEGFP233 amino acid

MKIEEGKLVIWINGDKGYNGLAEVGKKFEKDTGIKVTVEHPDKLEEKFPQVAATGDGPDIIFWAHDRFG

GYAQSGLLAEITPDKAFQDKLYPFTWDAVRYNGKLIAYPIAVEALSLIYNKDLLPNPPKTWEEIPALDK

ELKAKGKSALMFNLQEPYFTWPLIAADGGYAFKYENGKYDIKDVGVDNAGAKAGLTFLVDLIKNKHMNA

DTDYSIAEAAFNKGETAMTINGPWAWSNASYNVFIMADKQKNGIKANFKIRHNIEDGGVQLAYHYQQNT

PIGDGPVLLPDNHYLSVQSKLSKDPNEKRDHMVLLEFVTAAGITLGMDELYKGGTGGSMVSKGEELFTG

VVPILVELDGDVNGHKFSVSGEGEGDATYGKLTLKFICTTGKLPVPWPTLVTTLTYGVQCFSRYPDHMK

QHDFFKSAMPEGYIQERTIFFKDDGNYKTRAEVKFEGDTLVNRIELKGIDFKEDGNILGHKLEYNFNAS

SNIDTSKVNYGVTVLPTFKGQPSKPFVGVLSAGINAASPNKELAKEFLENYLLTDEGLEAVNKDKPLGA

VALKSYEEELAKDPRIAATMENAQKGEIMPNIPQMSAFWYAVRTAVINAASGRQTVDEALKDAQTRITK

*

>MBP_cpEGFP236 nucleotide

ATGAAAATCGAAGAAGGTAAACTGGTAATCTGGATTAACGGCGATAAAGGCTATAACGGACTCGCTGAA

GTCGGTAAGAAATTCGAGAAAGATACCGGAATTAAAGTCACCGTTGAGCATCCGGATAAACTGGAAGAG

AAATTCCCACAGGTTGCGGCAACTGGCGATGGCCCTGACATTATCTTCTGGGCACACGACCGCTTTGGT

GGCTACGCTCAATCTGGCCTGTTGGCTGAAATCACCCCGGACAAAGCGTTCCAGGACAAGCTGTATCCG

TTTACCTGGGATGCCGTACGTTACAACGGCAAGCTGATTGCTTACCCGATCGCTGTTGAAGCGTTATCG

CTGATTTATAACAAAGATCTGCTGCCGAACCCGCCAAAAACCTGGGAAGAGATCCCGGCGCTGGATAAA

GAACTGAAAGCGAAAGGTAAGAGCGCGCTGATGTTCAACCTGCAAGAACCGTACTTCACCTGGCCGCTG

ATTGCTGCTGACGGGGGTTATGCGTTCAAGTATGAAAACGGCAAGTACGACATTAAAGACGTGGGCGTG

GATAACGCTGGCGCGAAAGCGGGTCTGACCTTCCTGGTTGACCTGATTAAAAACAAACACATGAATGCA

GACACCGATTACTCCATCGCAGAAGCTGCCTTTAATAAAGGCGAAACAGCGATGACCATCAACGGCCCG

TGGGCATGGTCCAACATCGACACCGCATCTTATAACGTCTTTATCATGGCCGACAAGCAGAAGAACGGC

ATCAAGGCGAACTTCAAGATCCGCCACAACATCGAGGACGGCGGCGTGCAGCTCGCCTATCACTACCAG

CAGAACACCCCCATCGGCGACGGCCCCGTGCTGCTGCCCGACAACCACTACCTGAGCGTGCAGTCCAAA

CTGAGCAAAGACCCCAACGAGAAGCGCGATCACATGGTCCTGCTGGAGTTCGTGACCGCCGCCGGGATC

ACTCTCGGCATGGACGAGCTGTACAAGGGCGGTACCGGAGGGAGCATGGTGAGCAAGGGCGAGGAGCTG

TTCACCGGGGTGGTGCCCATCCTGGTCGAGCTGGACGGCGACGTAAACGGCCACAAGTTCAGCGTGTCC

GGCGAGGGCGAGGGCGATGCCACCTACGGCAAGCTGACCCTGAAGTTCATCTGCACCACCGGCAAGCTG

CCCGTGCCCTGGCCCACCCTCGTGACCACCCTGACCTACGGCGTGCAGTGCTTCAGCCGCTACCCCGAC

CACATGAAGCAGCACGACTTCTTCAAGTCCGCCATGCCCGAAGGCTACATTCAGGAGCGCACCATCTTC

TTCAAGGACGACGGCAACTATAAGACACGCGCTGAGGTTAAGTTCGAGGGCGACACTCTGGTTAACCGC

ATCGAGCTGAAGGGCATCGACTTCAAGGAGGACGGCAACATCCTGGGCCATAAGCTTGAATATAACTTC

AACGCATCTGACACCAGCAAAGTGAATTATGGTGTAACGGTACTGCCGACCTTCAAGGGTCAACCATCC

AAACCGTTCGTTGGCGTGCTGAGCGCAGGTATTAACGCCGCCAGTCCGAACAAAGAGCTGGCGAAAGAG

TTCCTCGAAAACTATCTGCTGACTGATGAAGGTCTGGAAGCGGTTAATAAAGACAAACCGCTGGGTGCC

GTAGCGCTGAAGTCTTACGAGGAAGAGTTGGCGAAAGATCCACGTATTGCCGCCACCATGGAAAACGCC

CAGAAAGGTGAAATCATGCCGAACATCCCGCAGATGTCCGCTTTCTGGTATGCCGTGCGTACTGCGGTG

ATCAACGCCGCCAGCGGTCGTCAGACTGTCGATGAAGCCCTGAAAGACGCGCAGACTCGTATCACCAAG

TAA

>MBP_cpEGFP236 amino acid

MKIEEGKLVIWINGDKGYNGLAEVGKKFEKDTGIKVTVEHPDKLEEKFPQVAATGDGPDIIFWAHDRFG

GYAQSGLLAEITPDKAFQDKLYPFTWDAVRYNGKLIAYPIAVEALSLIYNKDLLPNPPKTWEEIPALDK

ELKAKGKSALMFNLQEPYFTWPLIAADGGYAFKYENGKYDIKDVGVDNAGAKAGLTFLVDLIKNKHMNA

DTDYSIAEAAFNKGETAMTINGPWAWSNIDTASYNVFIMADKQKNGIKANFKIRHNIEDGGVQLAYHYQ

QNTPIGDGPVLLPDNHYLSVQSKLSKDPNEKRDHMVLLEFVTAAGITLGMDELYKGGTGGSMVSKGEEL

FTGVVPILVELDGDVNGHKFSVSGEGEGDATYGKLTLKFICTTGKLPVPWPTLVTTLTYGVQCFSRYPD

HMKQHDFFKSAMPEGYIQERTIFFKDDGNYKTRAEVKFEGDTLVNRIELKGIDFKEDGNILGHKLEYNF

NASDTSKVNYGVTVLPTFKGQPSKPFVGVLSAGINAASPNKELAKEFLENYLLTDEGLEAVNKDKPLGA

VALKSYEEELAKDPRIAATMENAQKGEIMPNIPQMSAFWYAVRTAVINAASGRQTVDEALKDAQTRITK

*

>MBP_cpEGFP131 nucleotide

ATGAAAATCGAAGAAGGTAAACTGGTAATCTGGATTAACGGCGATAAAGGCTATAACGGACTCGCTGAA

GTCGGTAAGAAATTCGAGAAAGATACCGGAATTAAAGTCACCGTTGAGCATCCGGATAAACTGGAAGAG

AAATTCCCACAGGTTGCGGCAACTGGCGATGGCCCTGACATTATCTTCTGGGCACACGACCGCTTTGGT

GGCTACGCTCAATCTGGCCTGTTGGCTGAAATCACCCCGGACAAAGCGTTCCAGGACAAGCTGTATCCG

TTTACCTGGGATGCCGTACGTTACAACGGCAAGCTGATTGCTTACCCGATCGCTGTTGAAGCGTTATCG

CTGATTTATAACAAAGATCTGCTGCCGAACCCGCCAAAAACCTGGGAAGAGATCGCATCTTATAACGTC

TTTATCATGGCCGACAAGCAGAAGAACGGCATCAAGGCGAACTTCAAGATCCGCCACAACATCGAGGAC

GGCGGCGTGCAGCTCGCCTATCACTACCAGCAGAACACCCCCATCGGCGACGGCCCCGTGCTGCTGCCC

GACAACCACTACCTGAGCGTGCAGTCCAAACTGAGCAAAGACCCCAACGAGAAGCGCGATCACATGGTC

CTGCTGGAGTTCGTGACCGCCGCCGGGATCACTCTCGGCATGGACGAGCTGTACAAGGGCGGTACCGGA

GGGAGCATGGTGAGCAAGGGCGAGGAGCTGTTCACCGGGGTGGTGCCCATCCTGGTCGAGCTGGACGGC

GACGTAAACGGCCACAAGTTCAGCGTGTCCGGCGAGGGCGAGGGCGATGCCACCTACGGCAAGCTGACC

CTGAAGTTCATCTGCACCACCGGCAAGCTGCCCGTGCCCTGGCCCACCCTCGTGACCACCCTGACCTAC

GGCGTGCAGTGCTTCAGCCGCTACCCCGACCACATGAAGCAGCACGACTTCTTCAAGTCCGCCATGCCC

GAAGGCTACATTCAGGAGCGCACCATCTTCTTCAAGGACGACGGCAACTATAAGACACGCGCTGAGGTT

AAGTTCGAGGGCGACACTCTGGTTAACCGCATCGAGCTGAAGGGCATCGACTTCAAGGAGGACGGCAAC

ATCCTGGGCCATAAGCTTGAATATAACTTCAACGCATCTGAGATCCCGGCGCTGGATAAAGAACTGAAA

GCGAAAGGTAAGAGCGCGCTGATGTTCAACCTGCAAGAACCGTACTTCACCTGGCCGCTGATTGCTGCT

GACGGGGGTTATGCGTTCAAGTATGAAAACGGCAAGTACGACATTAAAGACGTGGGCGTGGATAACGCT

GGCGCGAAAGCGGGTCTGACCTTCCTGGTTGACCTGATTAAAAACAAACACATGAATGCAGACACCGAT

TACTCCATCGCAGAAGCTGCCTTTAATAAAGGCGAAACAGCGATGACCATCAACGGCCCGTGGGCATGG

TCCAACATCGACACCAGCAAAGTGAATTATGGTGTAACGGTACTGCCGACCTTCAAGGGTCAACCATCC

AAACCGTTCGTTGGCGTGCTGAGCGCAGGTATTAACGCCGCCAGTCCGAACAAAGAGCTGGCGAAAGAG

TTCCTCGAAAACTATCTGCTGACTGATGAAGGTCTGGAAGCGGTTAATAAAGACAAACCGCTGGGTGCC

GTAGCGCTGAAGTCTTACGAGGAAGAGTTGGCGAAAGATCCACGTATTGCCGCCACCATGGAAAACGCC

CAGAAAGGTGAAATCATGCCGAACATCCCGCAGATGTCCGCTTTCTGGTATGCCGTGCGTACTGCGGTG

ATCAACGCCGCCAGCGGTCGTCAGACTGTCGATGAAGCCCTGAAAGACGCGCAGACTCGTATCACCAAG

TAA

>MBP_cpEGFP131 amino acid

MKIEEGKLVIWINGDKGYNGLAEVGKKFEKDTGIKVTVEHPDKLEEKFPQVAATGDGPDIIFWAHDRFG

GYAQSGLLAEITPDKAFQDKLYPFTWDAVRYNGKLIAYPIAVEALSLIYNKDLLPNPPKTWEEIASYNV

FIMADKQKNGIKANFKIRHNIEDGGVQLAYHYQQNTPIGDGPVLLPDNHYLSVQSKLSKDPNEKRDHMV

LLEFVTAAGITLGMDELYKGGTGGSMVSKGEELFTGVVPILVELDGDVNGHKFSVSGEGEGDATYGKLT

LKFICTTGKLPVPWPTLVTTLTYGVQCFSRYPDHMKQHDFFKSAMPEGYIQERTIFFKDDGNYKTRAEV

KFEGDTLVNRIELKGIDFKEDGNILGHKLEYNFNASEIPALDKELKAKGKSALMFNLQEPYFTWPLIAA

DGGYAFKYENGKYDIKDVGVDNAGAKAGLTFLVDLIKNKHMNADTDYSIAEAAFNKGETAMTINGPWAW

SNIDTSKVNYGVTVLPTFKGQPSKPFVGVLSAGINAASPNKELAKEFLENYLLTDEGLEAVNKDKPLGA

VALKSYEEELAKDPRIAATMENAQKGEIMPNIPQMSAFWYAVRTAVINAASGRQTVDEALKDAQTRITK

*

>MBP_cpEGFP195 nucleotide

ATGAAAATCGAAGAAGGTAAACTGGTAATCTGGATTAACGGCGATAAAGGCTATAACGGACTCGCTGAA

GTCGGTAAGAAATTCGAGAAAGATACCGGAATTAAAGTCACCGTTGAGCATCCGGATAAACTGGAAGAG

AAATTCCCACAGGTTGCGGCAACTGGCGATGGCCCTGACATTATCTTCTGGGCACACGACCGCTTTGGT

GGCTACGCTCAATCTGGCCTGTTGGCTGAAATCACCCCGGACAAAGCGTTCCAGGACAAGCTGTATCCG

TTTACCTGGGATGCCGTACGTTACAACGGCAAGCTGATTGCTTACCCGATCGCTGTTGAAGCGTTATCG

CTGATTTATAACAAAGATCTGCTGCCGAACCCGCCAAAAACCTGGGAAGAGATCCCGGCGCTGGATAAA

GAACTGAAAGCGAAAGGTAAGAGCGCGCTGATGTTCAACCTGCAAGAACCGTACTTCACCTGGCCGCTG

ATTGCTGCTGACGGGGGTTATGCGTTCAAGTATGAAAACGGCAAGTACGACATTAAAGACGTGGGCGTG

GATAACGCTGGCGCGAAAGCGGGTCTGACCTTCCTGGTTGCATCTTATAACGTCTTTATCATGGCCGAC

AAGCAGAAGAACGGCATCAAGGCGAACTTCAAGATCCGCCACAACATCGAGGACGGCGGCGTGCAGCTC

GCCTATCACTACCAGCAGAACACCCCCATCGGCGACGGCCCCGTGCTGCTGCCCGACAACCACTACCTG

AGCGTGCAGTCCAAACTGAGCAAAGACCCCAACGAGAAGCGCGATCACATGGTCCTGCTGGAGTTCGTG

ACCGCCGCCGGGATCACTCTCGGCATGGACGAGCTGTACAAGGGCGGTACCGGAGGGAGCATGGTGAGC

AAGGGCGAGGAGCTGTTCACCGGGGTGGTGCCCATCCTGGTCGAGCTGGACGGCGACGTAAACGGCCAC

AAGTTCAGCGTGTCCGGCGAGGGCGAGGGCGATGCCACCTACGGCAAGCTGACCCTGAAGTTCATCTGC

ACCACCGGCAAGCTGCCCGTGCCCTGGCCCACCCTCGTGACCACCCTGACCTACGGCGTGCAGTGCTTC

AGCCGCTACCCCGACCACATGAAGCAGCACGACTTCTTCAAGTCCGCCATGCCCGAAGGCTACATTCAG

GAGCGCACCATCTTCTTCAAGGACGACGGCAACTATAAGACACGCGCTGAGGTTAAGTTCGAGGGCGAC

ACTCTGGTTAACCGCATCGAGCTGAAGGGCATCGACTTCAAGGAGGACGGCAACATCCTGGGCCATAAG

CTTGAATATAACTTCAACGCATCTCTGGTTGACCTGATTAAAAACAAACACATGAATGCAGACACCGAT

TACTCCATCGCAGAAGCTGCCTTTAATAAAGGCGAAACAGCGATGACCATCAACGGCCCGTGGGCATGG

TCCAACATCGACACCAGCAAAGTGAATTATGGTGTAACGGTACTGCCGACCTTCAAGGGTCAACCATCC

AAACCGTTCGTTGGCGTGCTGAGCGCAGGTATTAACGCCGCCAGTCCGAACAAAGAGCTGGCGAAAGAG

TTCCTCGAAAACTATCTGCTGACTGATGAAGGTCTGGAAGCGGTTAATAAAGACAAACCGCTGGGTGCC

GTAGCGCTGAAGTCTTACGAGGAAGAGTTGGCGAAAGATCCACGTATTGCCGCCACCATGGAAAACGCC

CAGAAAGGTGAAATCATGCCGAACATCCCGCAGATGTCCGCTTTCTGGTATGCCGTGCGTACTGCGGTG

ATCAACGCCGCCAGCGGTCGTCAGACTGTCGATGAAGCCCTGAAAGACGCGCAGACTCGTATCACCAAG

TAA

>MBP_cpEGFP195 amino acid

MKIEEGKLVIWINGDKGYNGLAEVGKKFEKDTGIKVTVEHPDKLEEKFPQVAATGDGPDIIFWAHDRFG

GYAQSGLLAEITPDKAFQDKLYPFTWDAVRYNGKLIAYPIAVEALSLIYNKDLLPNPPKTWEEIPALDK

ELKAKGKSALMFNLQEPYFTWPLIAADGGYAFKYENGKYDIKDVGVDNAGAKAGLTFLVASYNVFIMAD

KQKNGIKANFKIRHNIEDGGVQLAYHYQQNTPIGDGPVLLPDNHYLSVQSKLSKDPNEKRDHMVLLEFV

TAAGITLGMDELYKGGTGGSMVSKGEELFTGVVPILVELDGDVNGHKFSVSGEGEGDATYGKLTLKFIC

TTGKLPVPWPTLVTTLTYGVQCFSRYPDHMKQHDFFKSAMPEGYIQERTIFFKDDGNYKTRAEVKFEGD

TLVNRIELKGIDFKEDGNILGHKLEYNFNASLVDLIKNKHMNADTDYSIAEAAFNKGETAMTINGPWAW

SNIDTSKVNYGVTVLPTFKGQPSKPFVGVLSAGINAASPNKELAKEFLENYLLTDEGLEAVNKDKPLGA

VALKSYEEELAKDPRIAATMENAQKGEIMPNIPQMSAFWYAVRTAVINAASGRQTVDEALKDAQTRITK

*
